# Cortical representational geometry of diverse tasks reveals subject-specific and subject-invariant cognitive structures

**DOI:** 10.1101/2024.01.26.577334

**Authors:** Tomoya Nakai, Rieko Kubo, Shinji Nishimoto

## Abstract

The variability in brain function forms the basis for our uniqueness. Prior studies indicate smaller individual differences and larger inter-subject correlation (ISC) in sensorimotor areas than in the association cortex. These studies, deriving information from brain activity, leave individual differences in cognitive structures based on task similarity relations unexplored. This study quantitatively evaluates these differences by integrating ISC, representational similarity analysis, and vertex-wise encoding models using functional magnetic resonance imaging across 25 cognitive tasks. ISC based on cognitive structures enables subject identification with 100% accuracy using at least 14 tasks. ISC is larger in the fronto-parietal association and higher-order visual cortices, suggesting subject-invariant cognitive structures in these regions. Principal component analysis reveals different cognitive structure configurations within these regions. This study provides new evidence of individual variability and similarity in abstract cognitive structures.

## Introduction

What brain properties shape our uniqueness and commonality? Cognitive neuroscience researchers have often viewed inter-individual differences as random noise. However, an increasing number of studies have quantitatively examined such variability in brain structures and functions (Dubois and Adolphs 2016; Seghier and Price 2018; Michon et al. 2022). These studies reported distinct subject-specific activity patterns even after anatomically aligning individuals’ data into a template brain (Dubois and Adolphs 2016; Feilong et al. 2018), which are highly stable across different experimental sessions (Wang et al. 2015; Gordon et al. 2017; Gratton et al. 2018). Quantitative comparison of functional connectivity consistently found greater individual differences in the fronto-parietal association cortex, while smaller differences were observed in the early sensorimotor cortex (Mueller et al. 2013; Vanderwal et al. 2017; Seitzman et al. 2019; Cutts et al. 2022). Similar cortical organization of individual variability was observed in different developmental stages (Cui et al. 2020; Stoecklein et al. 2020), suggesting a possible link with the functional hierarchy along the sensorimotor-association cortical axis (Margulies et al. 2016; Keller et al. 2023).

Simultaneously, other studies assessed similarities in brain activity among individuals. A common methodology in these studies is inter-subject correlation (ISC) analysis, which calculates how temporal activity profiles or functional connectivity of different subjects synchronize when they are watching movies (Hasson et al. 2004, 2008), listening to speech and music (Wilson, Molnar-Szakacs, and Iacoboni 2008; Abrams et al. 2013; Simony et al. 2016), speech production (Silbert et al. 2014), and combining of multimodal stimuli (Setti et al. 2023). These studies have demonstrated that ISC is larger in the low-level sensory areas and smaller in the association cortices (Ren et al. 2017; Kauppi et al. 2017). As smaller individual differences in brain activity equate to similar brain activity patterns across individuals, this is likely an investigation of a similar phenomenon from a different angle.

While these approaches derive information directly from brain activity, they leave unclear how individuals differ in their abstract relational structures among diverse cognitive functions, referred to as *cognitive structure*. For instance, while some individuals may exhibit a strong link between language and visual functions, others may show a stronger association between language and auditory functions (**Fig. 1A**). To examine such cognitive structures, researchers have developed the representational similarity analysis (RSA) (Kriegeskorte and Kievit 2013). This technique models representations of discrete tasks or stimuli through multivoxel activity patterns, enabling computation of cognitive structures among diverse tasks (Nakai and Nishimoto 2020, 2022). So far, RSA has mainly been used to investigate shared activity patterns at the group level (Kriegeskorte, Goebel, and Bandettini 2006; Devereux et al. 2013; Peelen and Downing 2023) and to compare with behavioral patterns (Charest et al. 2014; Brooks et al. 2019; Finn et al. 2020), but little is known about RSA’s effectiveness as a basis for inter-subject variability and similarity in cognitive structures or about the brain regions that represent such variability.

**Figure 1.**
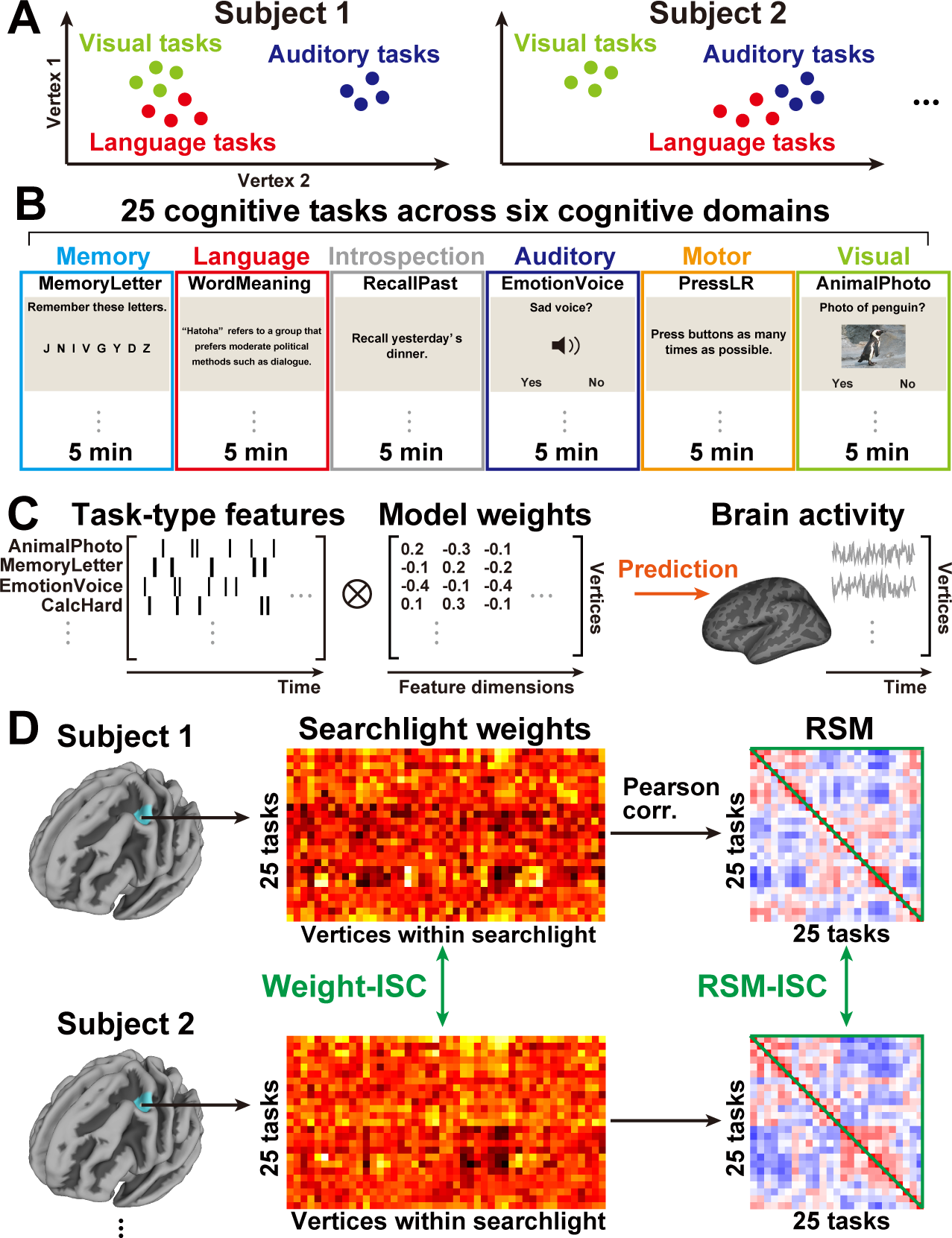
Evaluation of cognitive structures based on diverse tasks. **(A)** Schematic image of individual variability in cognitive structures. Tasks from three cognitive domains (visual, auditory, and language) are mapped onto the cortical representational space. **(B)** Subjects engaged in 25 cognitive tasks spanning six cognitive domains, with fMRI data collected during a 5-min duration for each cognitive domain. **(C)** Vertex-wise encoding models were constructed by representing brain activity through discrete task-type features, composed of a series of one-hot vectors. Weight matrices were computed using Ridge regression. (**D**) In the vicinity of each target cortical vertex, weight vectors were extracted from all vertices within a 9-mm geodesic distance (searchlight) on the cortical surface. The weight-based inter-subject correlation (weight-ISC) was determined by directly comparing the weight vectors of all subject pairs, averaged within the searchlight vertices. A representational similarity matrix (RSM) was generated for each searchlight by combining the weight vectors, and the RSM-ISC was computed by comparing the upper triangular components of RSMs for all subject pairs.

We address these challenges by integrating ISC and RSA across various cognitive tasks using vertex-wise encoding models (Naselaris et al. 2011). Encoding models quantitatively predict brain activity based on a combination of features from presented stimuli. This approach has been applied to model brain response patterns in visual (Kay, Naselaris, et al. 2008; Nishimoto et al. 2011; Çukur et al. 2013), auditory (Norman-Haignere, Kanwisher, and McDermott 2015; Nakai, Koide-Majima, and Nishimoto 2021), emotion (Horikawa et al. 2020; Koide-Majima, Nakai, and Nishimoto 2020), language (Huth et al. 2016; Nishida and Nishimoto 2018; Nakai, Yamaguchi, and Nishimoto 2021), and mathematical domains (Nakai and Nishimoto 2023a, 2023b). The weight matrices obtained during the encoding model construction reflect how target features are represented in the brain (Nakai and Nishimoto 2020; Koide-Majima, Nakai, and Nishimoto 2020), facilitating subsequent RSA and ISC analyses based on the model weights.

To comprehensively assess individuals’ cognitive structures across a broad range of abilities, we acquired functional magnetic resonance imaging (fMRI) data from 59 subjects performing 25 tasks spanning six cognitive domains (memory, language, introspection, auditory, motor, and visual) (**Fig. 1B**; detailed task descriptions in Supplementary Methods). These tasks were based on our prior studies with over 100 cognitive tasks, revealing a hierarchical organization within the six cognitive domains (Nakai and Nishimoto 2020, 2022). The current experimental design, a condensed version of these studies, aims to cover brain activity in most cortical areas within a 30 min timeframe per person. For each subject, we developed a vertex-wise encoding model utilizing categorical task-type features and derived weight values corresponding to the 25 tasks (**Fig. 1C**). Subsequently, we computed ISC using searchlight-based representational similarity matrices (RSMs) among the 25 tasks (**Fig. 1D**). To assess the methodology’s potential, we initially examined whether the encoding models and resultant RSMs effectively captured individual differences in cognitive structures. We then explored the brain organization of ISC concerning cognitive structures, visualizing these structures. Finally, we investigated how the brain organization of cognitive structures was influenced by different feature types using artificial neural network (ANN) models. This study presents novel evidence on the brain organization underlying individual variability and similarity in abstract cognitive structures.

## Results

### Encoding models capture individual variability in cognitive structures

Our initial inquiry revolves around the effectiveness of vertex-wise encoding models in capturing diverse cognitive structures among individuals. To address this, we trained a task-type encoding model for each subject using four out of the five runs and subsequently predicted brain activity in the left-out run. The performance of intra-subject prediction was first assessed. We observed that intra-subject encoding models robustly predicted activity across the entire cortex (prediction accuracy, mean ± SD, 0.153 ± 0.025; sign permutation test across subjects, 97.2% of vertices were significant; sign permutation test, corrected *p* < 0.05) (**Fig. 2A**). These significant vertices formed an inclusion mask for subsequent analyses.

**Figure 2.**
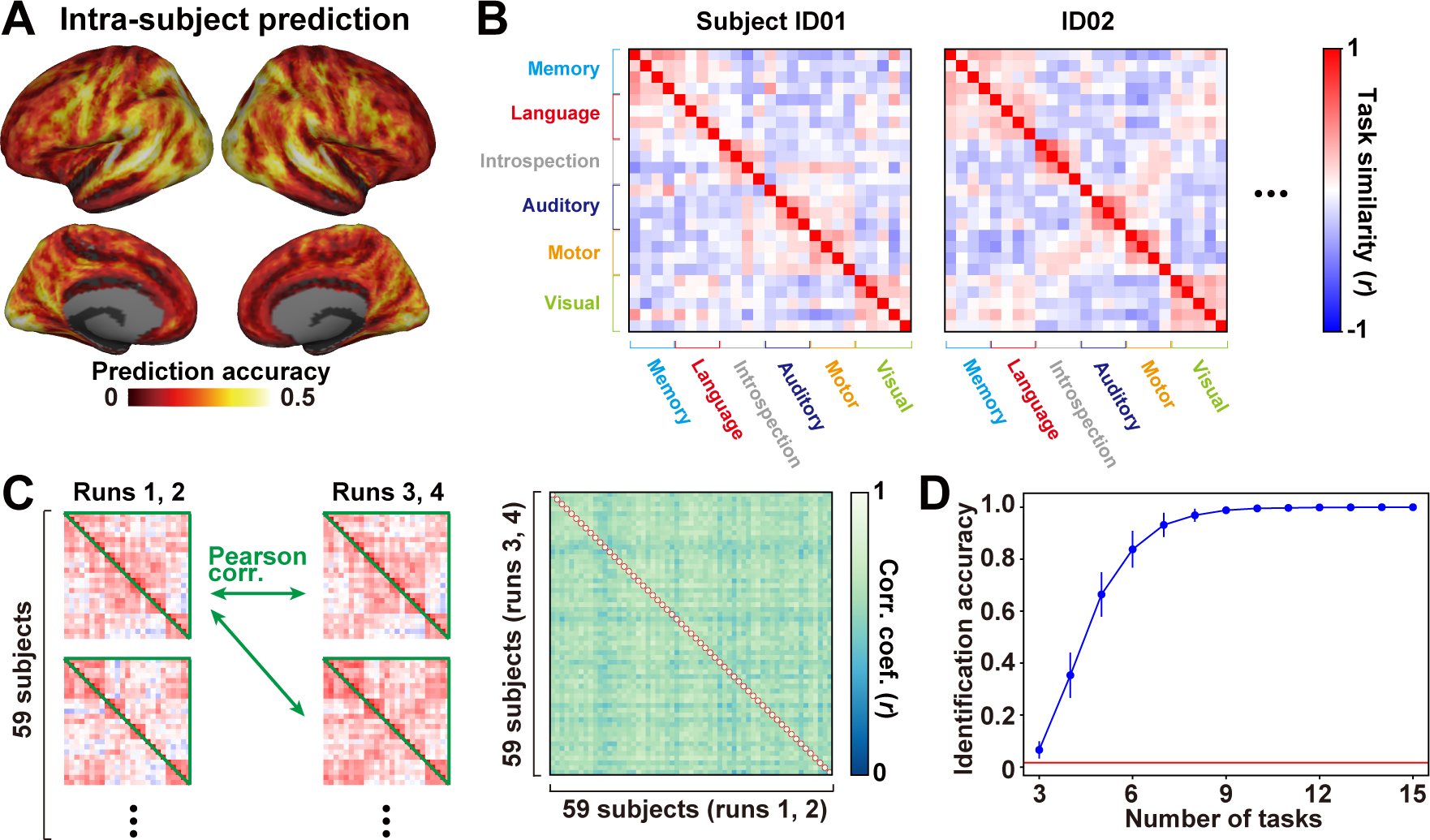
Encoding models capture subject-specific information. (**A**) The average prediction accuracy of the task-type encoding model is depicted on the standard cortical surface. The trained models are tested on the same subject (intra-subject prediction). **(B)** Example representational similarity matrices (RSMs) of two subjects calculated using all cortical vertices. Task pairs with a similar weight pattern are colored in red, while those with distinct weight patterns are in blue. **(C)** (left) Schematic representation of subject identification analysis. Two half-model RSMs are obtained using data from the 1^st^ and 2^nd^ runs for one half, and from the 3^rd^ and 4^th^ runs for the second half for each subject. Their upper triangular components are compared using Pearson’s correlation. (Right) The correlation coefficients of the half-model RSMs are shown for all subject pairs. For each row, the identified subject is marked with a red circle. **(D)** The modulation of identification accuracy is plotted as a function of the varying number of tasks. The red line indicates the chance-level accuracy (1/59).

We proceeded to assess the specificity of encoding models for individuals by applying these models to other individuals’ brain activity through *cross-subject prediction*. For each target subject, we averaged the prediction accuracy obtained using encoding models from the other 58 subjects. Once again, we observed significant prediction across the cortex (prediction accuracy, 0.054 ± 0.007; 94.6% of vertices were significant; sign permutation test, corrected *p* < 0.05) (**Fig. S1A**). Intra- and cross-subject prediction accuracies displayed a high correlation (Pearson’s correlation coefficient, *r* = 0.828). However, intra-subject prediction outperformed cross-subject prediction in 99.4% of cortical vertices (**Fig. S1B**). These results indicate that, although task-type encoding models can exhibit some degree of generalization to other individuals, they are highly specific to each individual.

To gauge individual differences in cognitive structures, we calculated an RSM among 25 tasks for each subject using weight matrices of all cortical vertices. RSMs were constructed based on Pearson’s correlation coefficients between all pairs of the 25 tasks. These RSMs demonstrated significant individual variability. For example, one subject (ID02) exhibited a substantial similarity (colored in red) between the visual and memory task domains, while another subject (ID01) displayed a lesser similarity (colored in blue) between these domains (**Fig. 2B**). The individual variability of task similarity (variance of similarity) was relatively large for the motor task domain, whereas it was smaller in the auditory and visual task domains (**Fig. S2**). For the other cognitive domains, the variance of similarity depended on the paired cognitive domains. These findings unequivocally demonstrate the existence of distinct cognitive structures among individuals.

We proceeded to assess whether these RSMs contained sufficient information for subject identification. For this purpose, we divided the original training data into two parts (runs 1–2 and runs 3–4) and reconstructed an encoding model for each part (**Fig. 2C**). RSMs were then calculated based on these half-models and compared across subjects. This analysis accurately identified all subjects (identification accuracy = 1.0; mean correlation coefficient, *r* = 0.928 ± 0.020 for the same subjects’ RSMs; 0.636 ± 0.035 for the different subjects’ RSMs).

Subject identification may be influenced by the task designs adopted in the experiment and the number of tasks included. To address this concern, we randomly selected a subset of the 25 tasks (ranging from 3 to 24 tasks) and calculated identification accuracy. The identification accuracy increased proportionally with the number of tasks included in the analysis and reached a plateau (i.e., identification accuracy = 1.0) at 14 tasks (**Fig. 2D**). This outcome suggests that our task set covers a much larger cognitive space than the minimal number of tasks required to identify subject-specific information.

### Subject-invariant cognitive structures are represented in higher-order brain regions

We endeavored to pinpoint the brain regions reflecting subject-invariant and subject-specific cognitive structures. In pursuit of this inquiry, we derived a weight matrix from the surface searchlight around each target vertex (within a 9 mm geodesic distance) and conducted ISC analysis based on the RSMs across 25 tasks (RSM-ISC, as depicted in **Figure 1C** right). Our observations unveiled relatively small RSM-ISC values around the central sulcus and superior temporal cortex, contrasting with large values in the fronto-parietal association cortices (**Fig. 3A**). While we utilized Pearson’s correlation coefficient as a similarity index in the preceding RSM calculation, aligning with ISC literature (Nastase et al. 2019), it’s noteworthy that other studies have employed Spearman’s correlation for comparing different RSMs (Finn et al. 2020; Ito and Murray 2022). To fortify our findings, we performed the same analysis using Spearman’s correlation, yielding similar results (**Fig. S3**), underscoring the resilience of our conclusions across distinct similarity indices.

**Figure 3.**
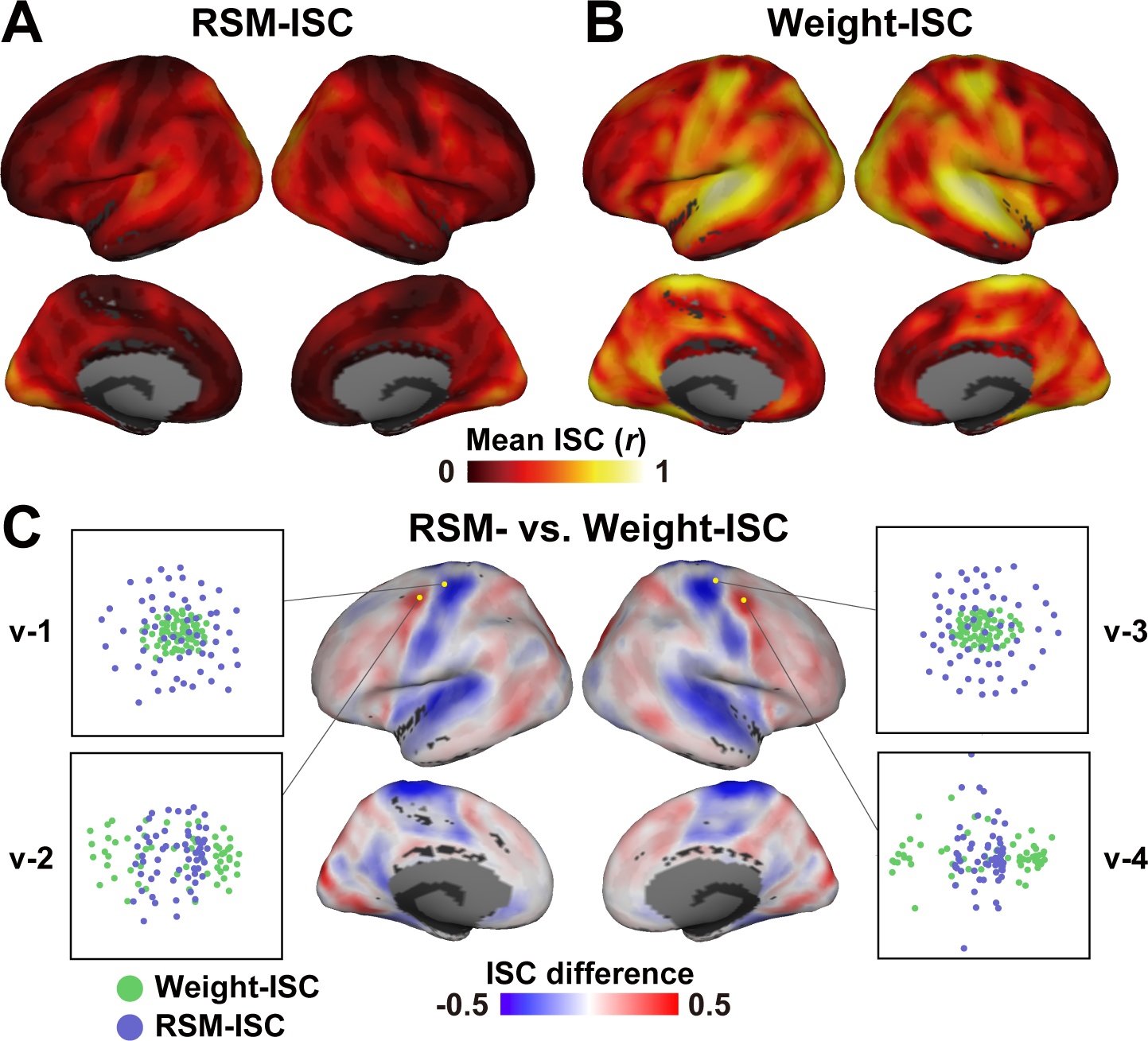
Larger RSM-ISC in higher-order brain regions. **(A)** Cortical surface mapping of Mean RSM-ISC and **(B)** Weight-ISC across all subject pairs. **(C)** Contrast between RSM-ISC and weight-ISC, where weight-ISC is corrected using the fitting curve obtained from simulation analysis and subtracted from RSM-ISC (**Fig. S3**). Vertices with higher RSM-ISC are colored in red, while those with higher (corrected) weight-ISC are in blue. Scatter plots depict multidimensional scaling analysis (MDS with IDIOSCAL scaling; see Methods for details) of weight-ISC and RSM-ISC values for data from four representative vertices across the 59 subjects.

The cognitive structures unveiled by the RSM-ISC encapsulate second-order information about the task weight matrix based on similarity relations. However, we did not assess whether the original weight matrices possessed the capacity to capture such cognitive structures. In pursuit of this, we calculated ISC by directly comparing model weights among subjects (weight-ISC, as depicted in **Figure 1C** center). In contrast to the RSM-ISC, we identified relatively large weight-ISC values around the central sulcus, as well as the superior temporal and occipital cortices (**Fig. 3B**). This dichotomy suggests that RSM-ISC and weight-ISC capture distinct facets of the brain’s representations of cognitive tasks.

Directly comparing weight-ISC and RSM-ISC might be inappropriate due to the possibility that some RSM-ISC components could arise from the arrangement of weight-ISC. Specifically, a scenario could exist where RSM-ISC has a large value if there is an array of synchronized vertices with large weight-ISCs spatially arranged around the target vertex (**Fig. S4,** left). Computational simulations indicated that RSM-ISC values tend to increase when weight-ISC values are elevated (**Fig. S4** right).

In consideration of this potential confounding factor, we corrected the weight-ISC value based on the fitting curve obtained from the simulation analysis and compared it with RSM-ISC (**Fig. 3C**). Our findings revealed that RSM-ISC was larger than the corrected weight-ISC in fronto-parietal association cortices, whereas the corrected weight-ISC was larger than RSM-ISC around the central sulcus and superior temporal cortex. Analyzing the distribution of differences between RSM-ISC and weight-ISC using pre-existing cortical parcellation (Yeo’s 17 network; **Fig. S5, S6**) (Schaefer et al. 2018), we noted that fronto-parietal networks (DorsAttnA, DorsAttnB, ContA, ContB, ContC) and the higher-order visual network (VisPeri) exhibited a larger RSM-ISC (difference between RSM- and weight-ISC, mean ± SD, 0.030 ± 0.028; standard deviation calculated across all subject pairs). In contrast, sensorimotor networks (TempPar, SomMotA, and SomMotB) demonstrated a larger weight-ISC (difference between RSM- and weight-ISC, −0.165 ± 0.037). Multidimensional scaling (MDS) analysis further visualized how RSMs and weight matrices differed across subjects (**Fig. 3C**). In the vertices (blue) where weight-ISC was larger than RSM-ISC (v-1, v-3), subjects were more closely located, indicating similarity in their representations using the weight-ISC value. Conversely, in the vertices (red) where RSM-ISC was larger than weight-ISC (v-2, v-4), subjects were more closely located using the RSM-ISC value. In summary, these findings indicate that, contrary to the previous literature on activity-based ISC, subject-invariant cognitive structures are organized in fronto-parietal and higher-order visual cortices.

### Distinct cognitive structures are represented on the cortical surface

The preceding analyses demonstrated the presence of subject-invariant cognitive structures in the fronto-parietal and higher-order visual areas. However, it remained uncertain whether such structures differed within these areas. To investigate this possibility, we conducted a principal component analysis (PCA). For data independence, we initially obtained a cortical mask of the RSM-vs. weight-ISC contrast (with significant vertices; sign permutation, corrected *p* < 0.05) using half of the subjects (29 subjects; ID01-ID29) and performed PCA with the remaining 30 subjects (ID30-ID59). We visualized distinct organization of cognitive structures across these areas in red, blue, and green colors corresponding to the top three principal components (PCs) (**Figs. 4A, S7**; also see results with the other half of subjects in **Figure S8**). These three PCs explained 50.5% of the variances in the original weight matrices (**Fig. S7**).

**Figure 4.**
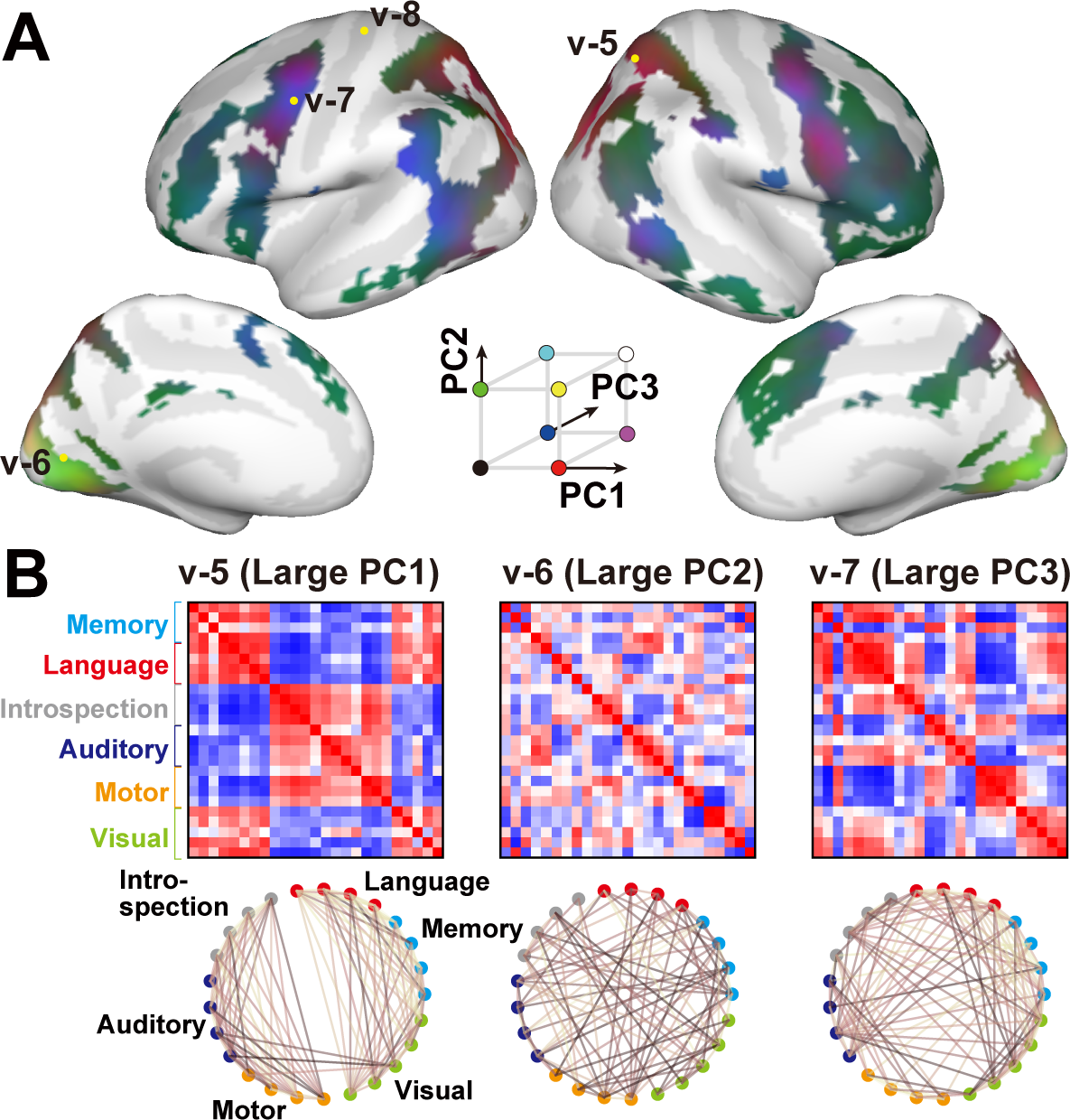
Cortical organization of subject-invariant cognitive structures. **(A)** A cortical map of distinct RSM patterns is revealed through principal component analysis (PCA). Mask vertices with a significant RSM-vs. weight-ISC value were derived from half of the subjects, and PCA was applied to the remaining subjects within the mask. Contributions of the top three PCs (PC1–PC3) are shown in red, green, and blue, respectively. **(B)** RSMs of three exemplary vertices (v-5, v-6, v-7) selected from **(A)**, along with their network visualization. RSMs are averaged across subjects. For clarity, only task pairs with a correlation coefficient (*r)* > 0.2 are connected.

Example vertices from each of the red, blue, and green regions clearly exhibited distinct cognitive structures. In a vertex from the red region (v-5, with a large PC1), tasks were organized separately based on their cognitive domains. Language, memory, and visual tasks constituted one cluster, while introspection, auditory, and motor tasks formed another distinct cluster (average RSM across subjects; **Fig. 4B**, left). In a vertex from the green region (v-6, with a large PC2), tasks were more tightly connected to tasks of different domains and did not form recognizable clusters (**Fig. 4B**, center). A vertex from the blue region (v-7, with a large PC3) showed an intermediate organization between the red and green vertices (**Fig. 4B**, right). Auditory and introspection tasks were connected to language, memory, and visual tasks, but motor tasks were not fully connected with these domains. In terms of individual RSMs, subjects displayed similar structures in the vertex with a large RSM-ISC (v-5, mean RSM-ISC = 0.45; **Fig. S9A**), while no clear structure was found in the vertex with a small RSM-ISC (v-8, RSM-ISC = 0.08; **Fig. S9B**).

### Similar but narrower distributions are obtained using other latent features

The categorical model used in the preceding sections assumes distinct representations for different tasks, modeled by one-hot vectors. While this model-based approach provides valuable insights, it is not the exclusive method for analyzing brain activity induced by cognitive tasks. Specifically, conventional model-free ISC (response-ISC) relies on direct comparisons of brain response patterns between individuals (Nastase et al. 2019). The response-ISC exhibited a similar pattern to the weight-ISC (*r* = 0.927; **Fig. S10A, B**), indicating relatively large values in the sensorimotor areas. In contrast, the response-ISC showed less consistency with the RSM-ISC (*r* = 0.725; **Fig. S10C**), likely due to deviations in the fronto-parietal and higher-order visual cortices. These results suggest that subject-invariant cognitive structures can be revealed through model-based RSM-ISC analysis, distinct from conventional ISC analysis.

Within the model-based approach, alternative models incorporating latent visual and language features could be considered. All tasks, even auditory and motor tasks, contained visual information in the form of instructional text. Recent advances in ANNs allow the extraction of latent features from stimuli, enabling the decomposition of categorical tasks based on these latent variables (Nakai and Nishimoto 2023b). Consequently, we extracted latent visual features using a visual transformer (ViT) and language features using GPT-Neox from presented images and instruction texts, respectively, constructing additional encoding models (**Fig. 5A**). It is important to note that the goal of this analysis was not to identify the optimal model for cognitive tasks, and the two types of features selected here serve as examples of latent features. These encoding models significantly predicted vertices across the cortex for both intra-subject prediction (ViT; prediction accuracy = 0.139 ± 0.023; 96.2% of vertices were significant; GPT-Neox; prediction accuracy = 0.148 ± 0.025; 97.2% of vertices were significant; sign permutation test, corrected *p* < 0.05) and cross-subject prediction (ViT; prediction accuracy = 0.048 ± 0.007; 93.4% of vertices were significant; GPT-Neox; prediction accuracy = 0.050 ± 0.007; 94.6% of vertices were significant) (**Fig. S11**). These findings suggest that both visual and language latent features can capture brain activity patterns associated with the diverse cognitive tasks used in the present study.

**Figure 5.**
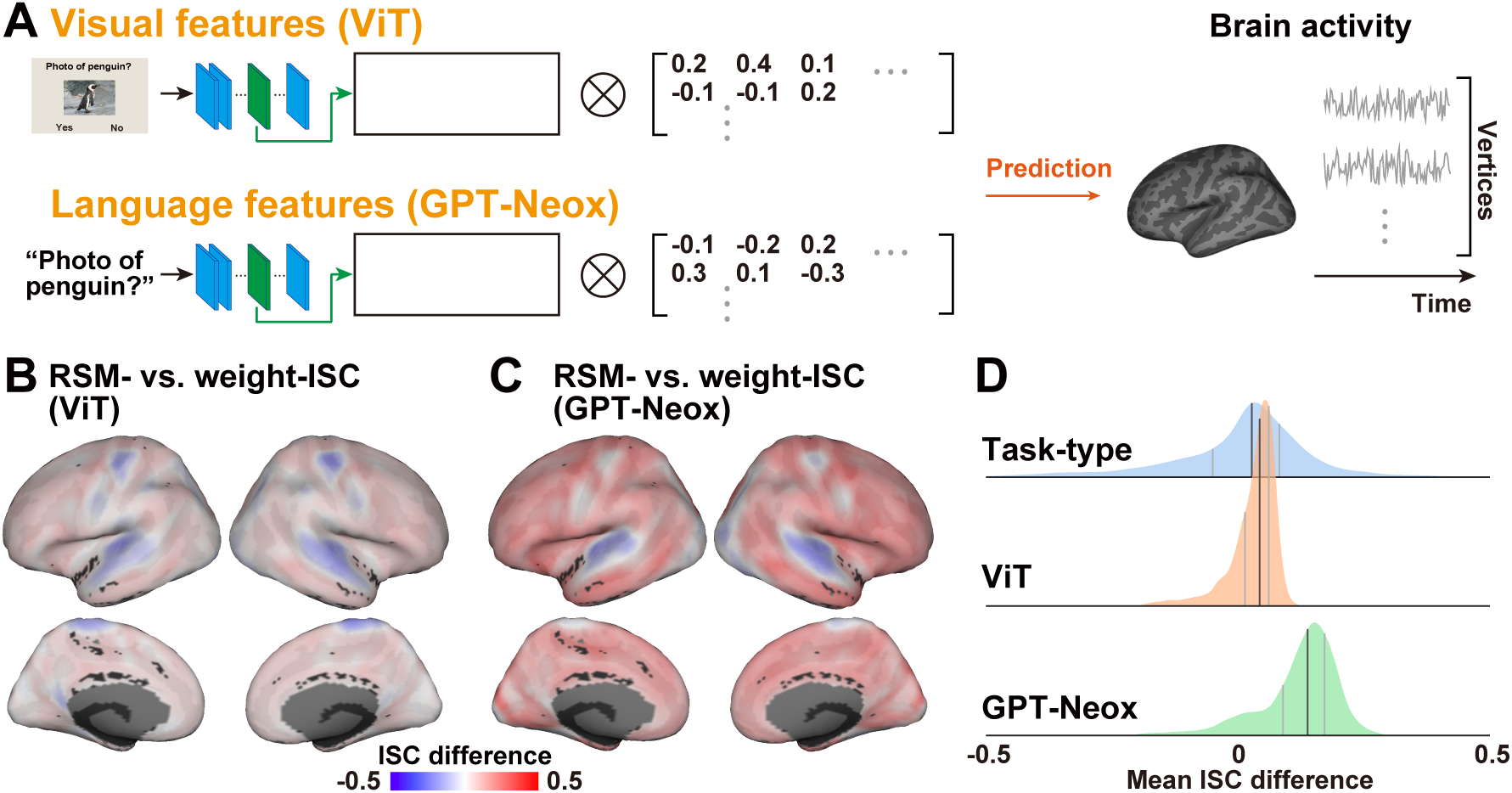
Latent neural network features exhibit similar but narrower distributions. **(A)** Illustration of encoding models utilizing visual (ViT) and language (GPT-Neox) neural network features. (**B-C**) Contrasts between RSM-ISC and weight-ISC for **(B)** visual features (ViT) and **(C)** language neural network features (GPT-Neox). **(D)** Distributions of the difference between RSM-ISC and weight-ISC across the cortex for the three types of features (task-type, ViT, and GPT-Neox).

Finally, we investigated whether the brain organization of cognitive structures is influenced by the latent features employed. Similar to the task-type features, we observed larger RSM-ISC values compared to weight-ISC values in the fronto-parietal networks (DorsAttnB, ContA, ContB, ContC) and the higher-order visual network (VisPeri) (difference between RSM- and weight-ISC; ViT, 0.020 ± 0.011; GPT-Neox, 0.113 ± 0.016) for both visual and language models (**Figs. 5B, 5C, S12**). Additionally, we identified smaller RSM-ISC values in the sensorimotor networks (TempPar, SomMotA, SomMotB) (difference between RSM- and weight-ISC; ViT, −0.028 ± 0.022; GPT-Neox, 0.057 ± 0.025). Furthermore, we observed an additional contribution of the limbic network (LimbicA) and salient ventral attention networks (SalVentAttnA and SalVentAttnB) to the RSM-ISC. The distribution of the ISC difference (i.e., RSM-vs. weight-ISC) was narrower than that of the task-type model (standard deviation; task-type model, 0.141; ViT model, 0.047; GPT-Neox, 0.078; **Fig. 5D**). Meanwhile, the encoding model based on the language features (GPT-Neox) yielded a distribution more on the positive side, with larger RSM-ISC values. These results suggest that, while the proposed methods demonstrate robustness, their patterns can vary depending on the properties of the selected features. Notably, a narrower distribution suggests that the distinctions between weight-ISC and RSM-ISC are less pronounced when using latent ANN features.

## Discussion

We have successfully demonstrated the effectiveness of our proposed method, RSM-ISC, in quantitatively assessing individual variability in cognitive structures. In this context, “cognitive structure” refers to a collection of representational similarities among 25 cognitive tasks, indicating whether pairs of tasks exhibit similar multivoxel activation patterns. In contrast, weight-ISCs, which involve the direct comparison of multivoxel patterns for each task across subjects, lack information regarding the relational aspects of different tasks. Moreover, while weight-ISC is confined to the spatial arrangement of brain activity patterns, RSM-ISC captures individual differences in the representational space based on similarity measures, presenting information abstracted from anatomical structures and activity patterns. The observed differences in cortical patterns between RSM-ISC and weight-ISC can be attributed to the impact of such constraints.

This approach stands apart from conventional ISC literature, which typically involves the direct comparison of temporal dynamics of brain activity between subjects during activities like movie viewing or story listening (Hasson et al. 2004, 2008; Simony et al. 2016; Nastase et al. 2019), yielding outcomes akin to weight-ISC. In the categorical task-type model, where cognitive tasks are treated as independent elements, model weights alone may not extract information beyond model-free brain activity. Although some studies have explored ISC based on spatial multivoxel patterns of brain activity (Chen et al. 2017; Sheng et al. 2023) and the similarity of multivoxel patterns between different time points (Guntupalli et al. 2016; Guntupalli, Feilong, and Haxby 2018), our method differs by not directly comparing activity patterns of cognitive tasks but rather focusing on the relations between model weights.

The inclusion of diverse tasks in our experimental design was crucial for implementing RSM-ISC. Without such tasks, it would be challenging to assess whether individuals’ cognitive structures are similar or different, particularly concerning functions like language and auditory processing. Unlike our previous studies (Nakai and Nishimoto 2020, 2022), where a small group performed 103 tasks over a 3-day MRI experiments, the current study, involving 59 subjects performing 25 tasks in a 30 min session, efficiently allowed us to capture cognitive structures across a larger participant pool. Despite the simplicity of the design, our encoding model effectively predicted over 90% of cortical vertices, indicating that the selected 25 tasks sufficiently covered diverse activity patterns in the cerebral cortex. Nevertheless, we observed that subjects can be accurately identified with at least 14 tasks, indicating the potential for optimizing the task set to more efficiently estimate cognitive structures in a shorter timeframe.

The searchlight analysis employing RSM-ISC revealed cortical patterns that contrast with findings from previous studies on individual variability and similarity (Mueller et al. 2013; Vanderwal et al. 2017; Ren et al. 2017; Kauppi et al. 2017). Notably, in our study, RSM-ISC exhibited larger values in the fronto-parietal networks (DorsAttnA, DorsAttnB, ContA, ContB, ContC) and the higher-order visual network (VisPeri), while showing smaller values in the low-level sensorimotor networks (TempPar, SomMotA, and SomMotB). Several studies have associated the fronto-parietal control network with general cognitive abilities, such as executive functions (Niendam et al. 2012; Friedman and Robbins 2022). This network has been considered a “hub” connecting various cognitive functions and facilitating task switching (Cole et al. 2013; Power et al. 2013; Ito and Murray 2022), suggesting systematic connections to diverse cognitive tasks across individuals. Although the higher-order visual network (VisPeri) may not traditionally be seen as a cognitive hub, a recent study suggests its involvement in the integration of multimodal information (Popham et al. 2021). Particularly in the current study, where task instructions were presented in text, the integration of visual and linguistic information may play a crucial role in processing cognitive tasks.

Further insights into the distinct roles of these networks were gained through PCA. A representative vertex from the fronto-parietal network exhibited discrete task clusters, while another vertex from the higher-order visual network showed intertwined relations. These results underscore the heterogeneity in cognitive structures even within subject-invariant networks. Notably, in the higher visual network, tasks within the same cluster often exhibited considerable dissimilarity in representational similarity. In contrast, most tasks within the same cluster demonstrated a similar representation in the fronto-parietal network. This suggests that in the latter network, cognitive tasks may be represented in a compressed form based on higher-order functional units or clusters, transcending superficial differences in visual stimuli, presentation time, and textual information.

The exploration of additional encoding models revealed that the model-based approach extended beyond discrete task-type features to include other latent features. The use of latent features from intermediate ANN layers has been prevalent in studies investigating the correspondence between ANNs and the brain (Horikawa and Kamitani 2017; Schrimpf et al. 2021; Nakai and Nishimoto 2023b). Given that latent features can be extracted from complex naturalistic stimuli like audio or movies, our approach holds potential for broader applicablity to various task-fMRI datasets. It could reveal individual differences not only in general cognitive structures but also in specific cognitive function structures, such as those related to object perception. Additionally, the RSM-ISC may offer a means to explore the extent to which latent features contain information about cognitive structures. The narrower distributions observed in both visual and language features might arise from shared visual and language components across the 25 tasks, making second-order information (i.e., RSM) less distinguishable from first-order information (i.e., model weights).

However, there are several limitations to the current study. First, while the 25 tasks adequately explained the activity of most cortical areas, this task set did not encompass certain crucial cognitive factors like odor perception or social communication. Components of cognitive structures related to such tasks may be absent from our analysis. Nevertheless, our method’s advantage lies in its applicability regardless of task type and model, enabling the investigation of individual variability in a wide range of cognitive components once relevant task-fMRI data is available.

Second, without the subtraction of weight-ISC, the raw RSM-ISC itself may not effectively distinguish cognitive structures in different cortical regions. As demonstrated in the simulation analysis, a group of vertices with similar model weight patterns across subjects may spontaneously generate high RSM-ISC values. The raw RSM-ISC includes the weight-ISC component as a confounder, obstructing the accurate quantification of individual differences in cognitive structures in their original form. To enhance the robustness of our results, further methodological development is necessary to separate cognitive structures from other components.

Finally, there may be a more suitable metric to assess cognitive structures than RSMs. RSM captures relational information based on the similarity of multivoxel patterns (Kriegeskorte and Kievit 2013). While widely utilized to test cognitive theories (Peelen and Downing 2023), this technique assumes that cognitive information is represented as spatial activity patterns akin to population coding. It overlooks the potential existence of other information formats, such as temporal dynamics and connectivity profiles. The low temporal resolution of fMRI data poses a limitation for examining the effects of temporal dynamics, and it remains to be determined whether the current methodology is applicable to other types of neuroimaging data with higher temporal resolution.

## Conclusions

The present study introduces a novel methodology for assessing individual similarity and variability in abstract cognitive structures by incorporating RSA and ISC. The findings highlight the involvement of higher-order cortical regions in shaping common cognitive structures across individuals. This research offers a fresh perspective on the neural basis of individual differences, highlighting the necessity for future research utilizing the present methodology to elucidate the origins of the disparities observed in comparison to prior investigations.

## Methods

### Subjects

Fifty-nine subjects, aged 20–55 years (19 females, all with normal vision), denoted as ID01– ID59, participated in this study. Written informed consent was obtained from all subjects before their participation, and the study received approval from the ethics and safety committee of the National Institute of Information and Communications Technology in Osaka, Japan.

### Stimuli and procedures

We designed 25 naturalistic tasks that required no pre-experimental training (see Supplementary Methods for detailed task descriptions). These tasks, derived from our prior studies (Nakai and Nishimoto 2020, 2022), were crafted to encompass a broad spectrum of cognitive domains (memory, language, introspection, auditory, motor, visual) within a 30 min experimental session. Each task comprised 10 instances, with 8 instances used in the training runs and 2 in the test run. Stimuli were presented on a projector screen inside the scanner (21.0 × 15.8° of visual angle at 30 Hz), and the root-mean-square of auditory stimuli was normalized. Subjects, equipped with MR-compatible ear tips. Presentation software (Neurobehavioral Systems, Albany, CA, USA) was used to control the stimulus presentation and the collection of behavioral data. To measure button responses, optic response pads with two buttons in each hand were used (HHSC-2 × 2, Current Designs, Philadelphia, PA, USA).

The experiment comprised 10 runs, with half dedicated to task-based fMRI (task-fMRI) and the remaining half to resting-state fMRI (rest-fMRI). The rest-fMRI data was not analyzed in the current study. Each task-fMRI run included 50 trials (2 trials for each of the 25 tasks), lasting 6–10 s per trial. Additionally, 10 s of scanning without a task was inserted at the beginning and end of each run; the former was excluded from the analysis. Both task-fMRI and rest-fMRI runs had a duration of 370 s. The task order was pseudorandomized, considering dependencies between certain tasks, such as the “MemoryLetter” and “MatchLetter” tasks, which were presented close to each other in time. The run order was counter-balanced across subjects. During task-fMRI runs, subjects received instructions on how to perform each task through instructional text presented as part of the stimuli (see **Fig. 1B**). Subjects underwent a brief training session on button responses.

### MRI data acquisition

The experiment was conducted using a 3.0 T scanner (MAGNETOM Vida; Siemens, Erlangen, Germany) with a 64-channel head coil. We scanned 72 interleaved axial slices, 2-mm thick without a gap, parallel to the anterior and posterior commissure line, using a T2*-weighted gradient-echo multiband echo-planar imaging sequence [repetition time (TR) = 1,000 ms; echo time (TE) = 30 ms; flip angle (FA) = 62°; field of view (FOV) = 192 × 192 mm^2^; resolution = 2 × 2 mm^2^; multiband factor = 6]. We obtained 370 volumes in each run, each following 10 dummy volumes. As an anatomical reference, high-resolution T1-weighted images of the whole brain were acquired from all subjects with a magnetization-prepared rapid acquisition gradient-echo sequence (TR = 2,530 ms; TE = 3.26 ms; FA = 9°; FOV = 256 × 256 mm^2^; voxel size = 1 × 1 × 1 mm^3^).

### Preprocessing

The preprocessing of both functional and anatomical MRI data was carried out using fMRIPrep 21.0.2 (Esteban et al. 2019). A B0 fieldmap was derived from phase-drift maps, which were measured using two successive gradient-recalled echo acquisitions. The corresponding phase maps were unwrapped using prelude (FSL 6.0.5.1).

The T1-weighted image underwent a correction for intensity non-uniformity using ANTs 2.3.3 N4BiasFieldCorrection (Avants et al. 2011). Following this, the image was skull-stripped using a Nipype implementation of the antsBrainExtraction.sh workflow. Brain tissue segmentation was conducted using fast (FSL 6.0.5.1) (Zhang, Brady, and Smith 2001). Brain surfaces were reconstructed using recon-all (FreeSurfer 6.0.1) (Dale, Fischl, and Sereno 1999). Lastly, volume-based spatial normalization to the standard space (MNI152NLin6Asym) was achieved through nonlinear registration with antsRegistration (ANTs 2.3.3).

For the functional scans, a reference volume and its skull-stripped version were initially produced using the custom methodology of fMRIPrep. Prior to any spatiotemporal filtering, head-motion parameters, which include six corresponding rotation and translation parameters, were estimated in relation to the reference scan using mcflirt (FSL 6.0.5.1) (Jenkinson et al. 2002). All functional scans underwent slice-time correction to the middle slice and were resampled onto their native space by applying the transforms to correct for head-motion. The reference scan was subsequently co-registered with the T1w reference using bbregister (FreeSurfer) (Greve and Fischl 2009). Three global signals were extracted within the cerebrospinal fluid (CSF), white matter (WM), and whole-brain masks. The blood oxygenation level-dependent time-series were resampled into the standard spaces (MNI152NLin6Asym). All resamplings were conducted with a single interpolation step by composing all the relevant transformations (i.e., head-motion transform matrices, susceptibility distortion correction, and co-registrations to anatomical and output spaces). Surface resampling was carried out using mri_vol2surf (FreeSurfer).

The resulting functional data underwent high-pass filtering at 0.01 Hz and low-pass filtering at 0.15 Hz. Confounding factors, including six head-motion parameters, global signals from the CSF, WM, and whole-brain masks, as well as the derivative of each factor, were eliminated using linear regression implemented in Scikit-learn. The functional data were then standardized to have a zero mean and unit variance for each of the 18,715 cortical vertices in the fsaverage5 space, excluding non-cortical vertices.

### Task-type features

The task-type features consisted of 25 one-hot vectors. Each time bin was assigned a value of 1 or 0, indicating whether one of the 25 tasks was performed during that period.

### Latent visual features (ViT)

Latent visual features were extracted using the vision transformer model (ViT; https://huggingface.co/docs/transformers/model_doc/vit) (Dosovitskiy et al. 2020). For each time bin, the output from the sixth layer (out of twelve layers) was extracted as visual features when each stimulus image was used as input. The resulting series of features were further reduced to 100 dimensions using a PCA.

### Latent language features (GPT-Neox)

Latent language features were extracted using the Japanese version of the transformer-based large language model, GPT-Neox https://huggingface.co/docs/transformers/model_doc/gpt_neox_japanese) (Black et al. 2022). When the instruction text of each stimulus was used as input, the output from the sixteenth layer (out of a total of thirty-two layers) was extracted as language features. These resultant series of features were then further reduced to 100 dimensions using PCA.

### Vertex-wise encoding model fitting

In the encoding model, each vertex’s cortical activity was modeled using a finite impulse response model. This model captured the slow hemodynamic response and its association with neural activity (Nishimoto et al. 2011; Kay, David, et al. 2008). The feature matrix, denoted as **F_E_** and of dimensions [T × 6N], was constructed by concatenating sets of [T × N] feature matrices, with six temporal delays ranging from 2 to 7 s (where T = number of samples; N = number of features). The cortical response denoted as **R_E_** and of dimensions [T × V], was then modeled by multiplying the feature matrix **F_E_** by the weight matrix **W_E_** of dimensions [6N × V] (where V = number of vertices).

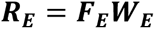

The weight matrix **W_E_** was obtained using L2-regularized linear regression with the training dataset, which consisted of 1,110 to 1,480 samples depending on the subjects. The optimal regularization parameter was determined using 5-fold cross-validation, where the regularization parameters varied from 1 to 10^6^ across seven different values.

The test dataset comprised 370 samples. The accuracy of the predictions was evaluated using Pearson’s correlation coefficient between the predicted and actual test samples. Statistical significance (one-sided) was calculated using a sign permutation test (*p* < 0.05), and multiple comparisons were corrected using the false-discovery rate (FDR) correction method (Benjamini and Hochberg 1995). Prediction accuracy was computed in both intra-subject and cross-subject manners. In the intra-subject prediction, the predicted samples and test samples were from the same subject. In the cross-subject prediction, the predicted samples and test samples were from different subjects. Significant cortical vertices in the inter-subject prediction were used as an inclusive mask in subsequent analyses. For visualizing data on cortical maps, we utilized pycortex (Gao et al. 2015).

### Whole cortex RSM

A whole-cortex RSM was obtained by calculating Pearson’s correlation coefficient (*r*) between all pairs of task weight vectors. These vectors contained data on all vertices, corresponding to the rows of the weight matrix, and were averaged for the six temporal delays. The resulting [25, 25] matrix shows how task pairs are similar or distinct (*r* = 1: perfectly similar; *r* = −1: perfectly distinct).

### Subject identification analysis

We conducted a subject identification analysis to assess whether RSMs contain enough information to differentiate between subjects. The original training data was initially split into two samples (comprising the 1^st^ and 2^nd^ runs and the 3^rd^ and 4^th^ runs). RSMs were computed for both samples, and their upper triangular matrices were compared using Pearson’s correlation coefficient. Specifically, correlation coefficients were calculated in both within-subject and cross-subject manners. A subject was deemed to be successfully identified if their within-subject correlation coefficient was larger than the cross-subject correlation coefficients between that subject and all other subjects.

We further repeated this analysis by incrementally increasing the number of tasks included in the RSMs from 3 to 24 (noting that we need at least three tasks to vectorize the upper triangular matrices). Target tasks were randomly selected from the original 25 tasks, and the mean identification accuracy across subjects was calculated. This process was repeated 100 times for each number of tasks.

### ISC analysis

To quantitatively assess the extent to which different subjects exhibit similar brain activity or representations, we employed three distinct methods for computing ISC: (1). Weight-ISC: ISC was directly computed from the weight matrices obtained from the vertex-wise encoding models. Weight vectors were extracted from vertices within a 9-mm geodesic distance around each target vertex on the cortical surface (i.e., a searchlight sphere). The resulting weight vectors, representing 25 task elements, were averaged across six temporal delays and within the target searchlight. Subsequently, ISC was computed at each target vertex by calculating Pearson’s correlation coefficient between the mean weight vectors of all subject pairs (a total of 1,711 pairs) and then averaging across subject pairs. (2) RSM-ISC individuals’ RSM was initially calculated by computing Pearson’s correlation coefficient between all pairs of task weight vectors extracted from the weight matrix within the target searchlight sphere. These weight vectors were not averaged across searchlight vertices, and the number of elements corresponded to the number of vertices. ISC was then computed based on the vectorized upper triangular part of the RSMs in a manner similar to weight-ISC. To ensure the robustness of the method, an additional analysis was performed using Spearman’s correlation coefficient as a similarity index. (3) Response-ISC brain response data from all training runs were concatenated and averaged across vertices within the target searchlight sphere. ISC was computed based on the concatenated response data for each target searchlight.

### ISC simulation

To investigate the relationship between weight-ISC and RSM-ISC, we conducted simulations of weight matrices. We created a reference weight matrix with 45 vertices ([25, 45] matrix) where the weight values adhered to a standard normal distribution. The number of vertices was determined based on the average number of searchlight vertices across the cortex. Using this reference weight matrix (denoted as *X_ref_*), we generated a dataset of 59 entries (matching the number of subjects in the current experiment) by varying the signal-to-noise ratio (SNR) value from 0 to 1 in increments of 0.05:

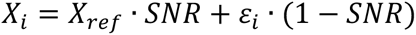

Where *ε_i_* represents a random noise matrix that follows a standard normal distribution, and *X_i_* denotes the simulated data for the *i*^th^ subject. We computed the weight-ISC and RSM-ISC using these *X_i_* s for each SNR value. The resulting simulated plot was then fitted with a quadratic polynomial function using a least-square fitting method.

### Multidimensional scaling

To visualize the representational relationships among subjects, we employed multidimensional scaling (MDS). We used the ISC value as a distance metric, also known as correlation distance. MDS can map different subjects onto a two-dimensional space based on their correlation distances, positioning subjects with high correlation coefficients closer together. However, one limitation of MDS is that maps from different sources (weight-ISC and RSM-ISC) cannot be directly compared. To address this issue, we adopted individual difference scaling (IDIOSCAL) (Carroll and Chang 1970). IDIOSCAL accounts for scaling variations due to different sources by using a weighted distance, thereby enabling the mapping of both weight-ISC and RSM-ISC of the 59 subjects onto the same two-dimensional space. This analysis was conducted using the SMACOF package in R (de Leeuw and Mair 2009).

### PCA

To explore the different cognitive structures on the cortical surface, we conducted a PCA based on the vectorized upper triangular part of the RSMs, which had _25_C_2_ = 300 dimensions. We only considered vertices that exhibited a significantly larger RSM-ISC than weight-ISC, with a significance level of *p* < 0.05 (determined by a sign permutation test with FDR correction). The vectorized RSMs were concatenated across all included vertices, and PCA was applied to the concatenated matrix. To visualize the cortical organization of the cognitive structures, we extracted and normalized the PCA scores from each vertex. The resulting cortical map indicated the relative contribution of each cortical vertex to the top three PCs.

The cognitive structure of the 25 tasks was further visualized as a network at each vertex based on their searchlight RSM, with a threshold of *r* = 0.2.

## Data and code availability

The source data and analysis code used in the current study are available upon reasonable request.

## Acknowledgements

We thank MEXT/JSPS KAKENHI (grant numbers JP24H02172 and JP24H01559 for T.N., JP15H05311 and JP18H05522 for S.N.), JST CREST JPMJCR18A5, ERATO JPMJER1801 (for S.N.), and H2020 Marie Skłodowska-Curie Actions (grant number 101023033 for T.N.) for partial financial support of this study. The funders had no role in the study design, data collection and analysis, decision to publish, or preparation of the manuscript.

## Author contributions

**T.N.**: Conceptualization, Methodology, Formal analysis, Visualization, Writing-Original draft preparation. **R.K.**: Data collection, Writing - Review & Editing. **S.N.**: Supervision, Writing - Review & Editing, Funding acquisition.

## Supplementary Information

**Figure S1.**
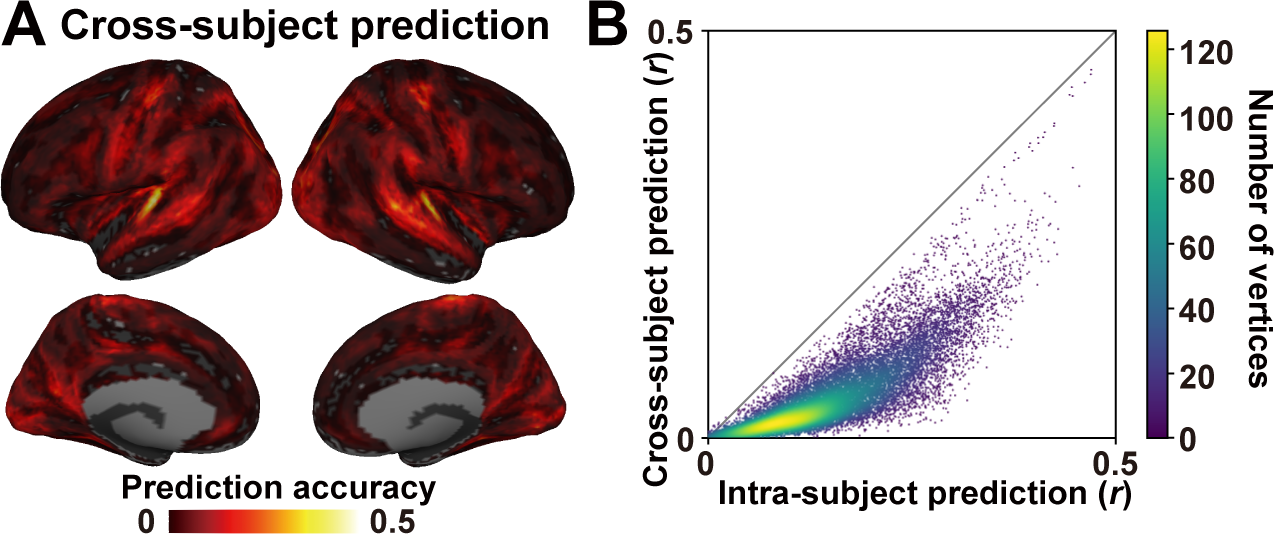
Cross-subject encoding model prediction. (**A**) The mean prediction accuracy of the task-type encoding model is represented on the standard cortical surface. Models are tested with different subjects from model training. Part **(B)** A scatter plot depicts the relationship between intra-subject and cross-subject prediction accuracy across the cortex.

**Figure S2.**
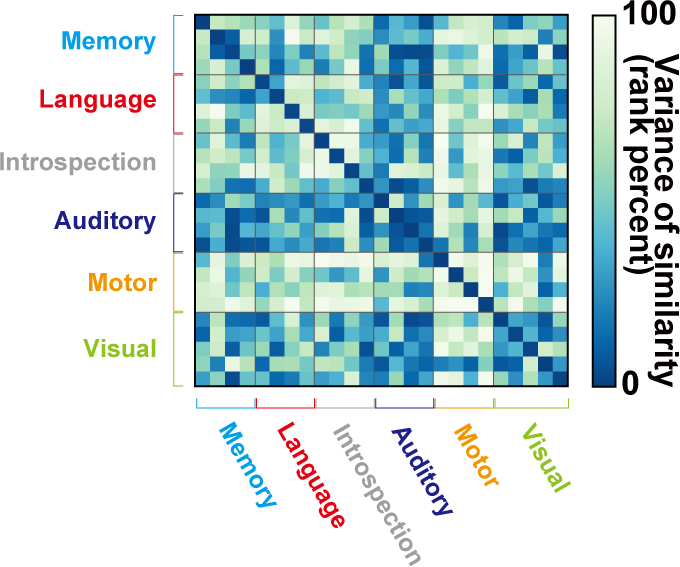
Variance of task similarity. The variance of similarity is determined by calculating the standard deviation of similarity values across all subjects. To enhance intelligibility, the variance values are converted to a rank percentile format.

**Figure S3.**
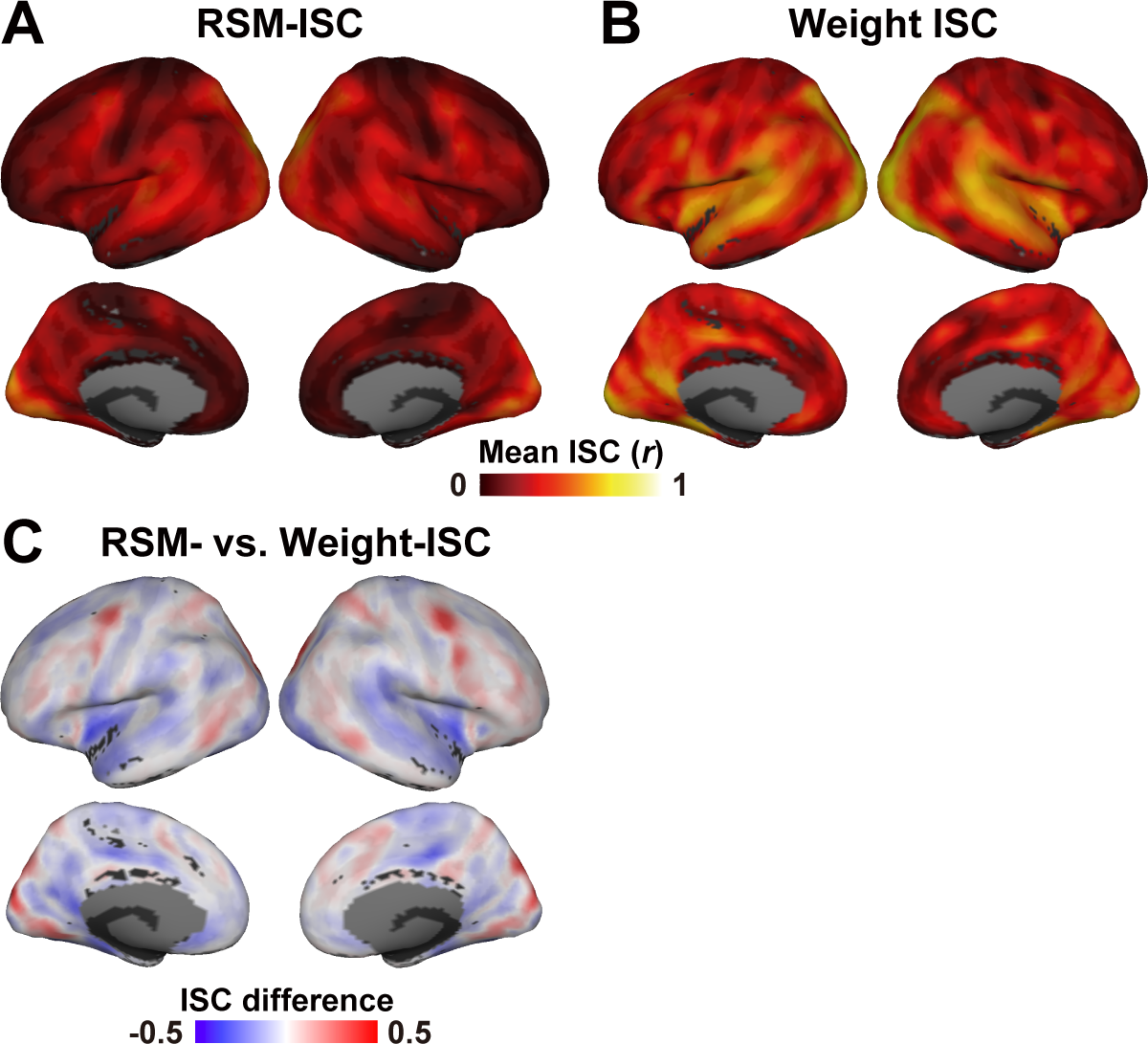
Inter-subject correlation (ISC) using Spearman’s correlation similarity. **(A)** Mean representational similarity matrix (RSM-) and weight-ISC and **(B)** ISC across all subject pairs are projected onto the cortical surface. **(C)** Weight-ISC is adjusted using the fitting curve derived from the simulation analysis and subtracted from the RSM-ISC. Vertices with a higher RSM-ISC are represented in red, whereas those with a higher (corrected) weight-ISC are depicted in blue.

**Figure S4.**
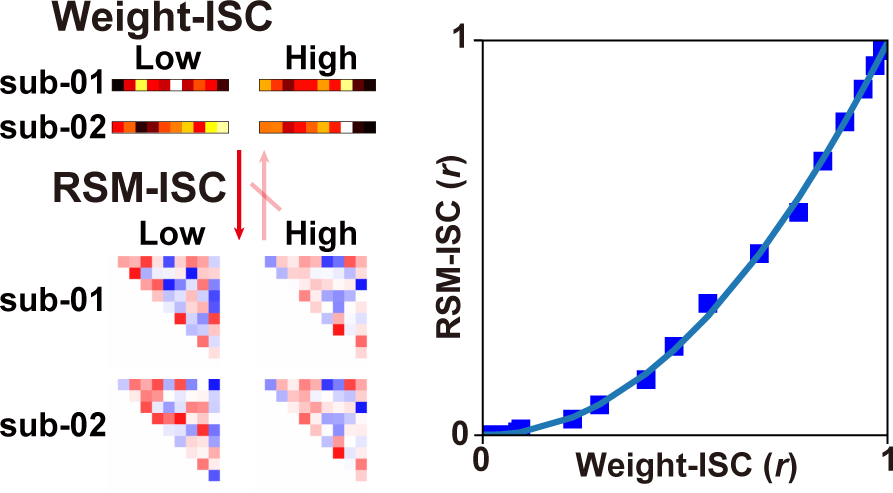
ISC simulation analysis. Schematic figure depicts the results of the simulation analysis. Low/high RSM-ISC values can emerge from corresponding low/high weight-ISC values. The simulated RSM-ISC is graphed as a function of weight-ISC, with an increasing signal-to-noise ratio. The plotted data is subsequently fitted with a quadratic polynomial function using least-squares fitting.

**Figure S5.**
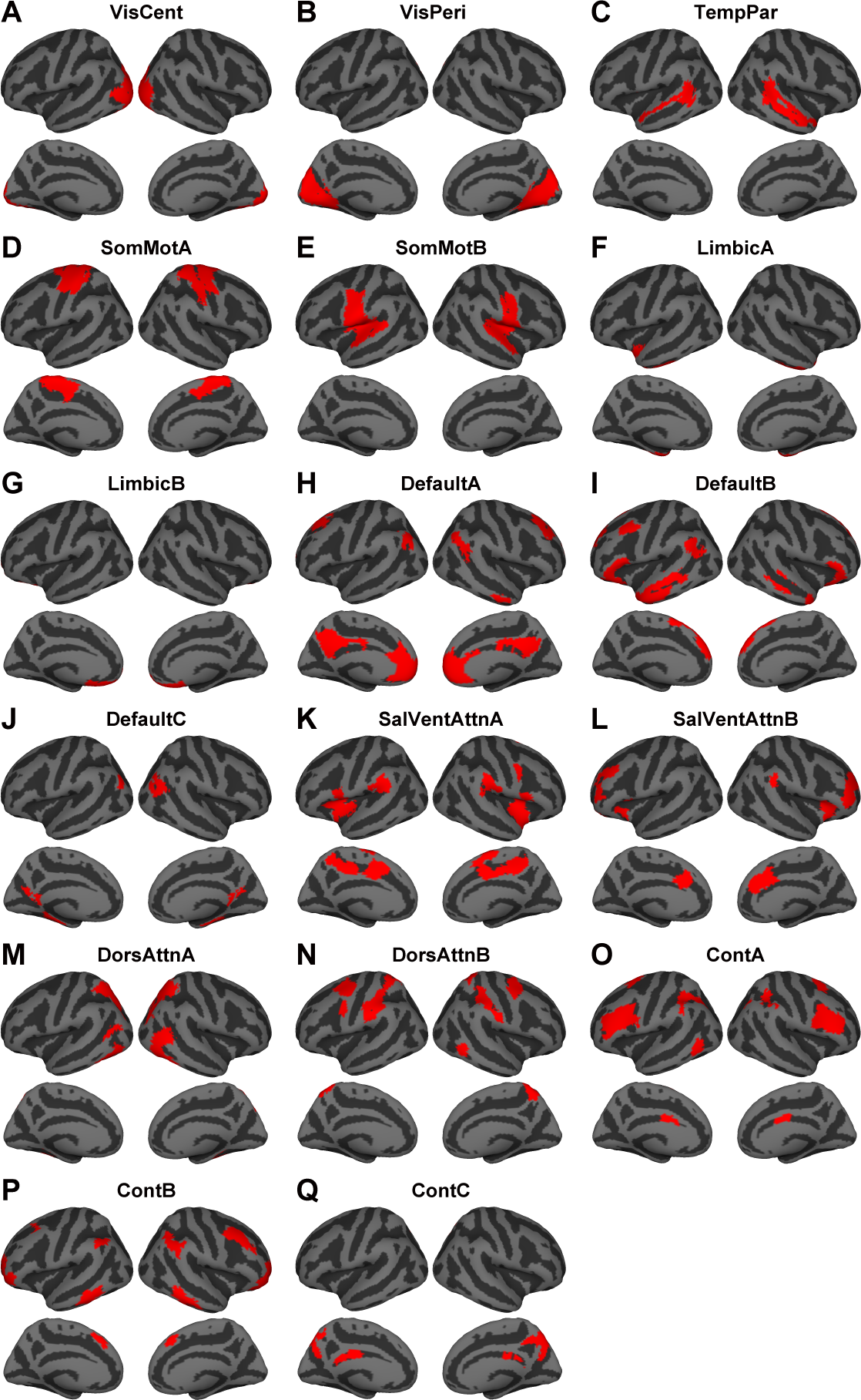
Schaefer’s 17 network atlas. Each network in this atlas was derived from 400 parcel parcellations as provided by Schaefer et al. (2018). These parcellations were then resampled and mapped onto the fsaverage5 cortical surface.

**Figure S6.**
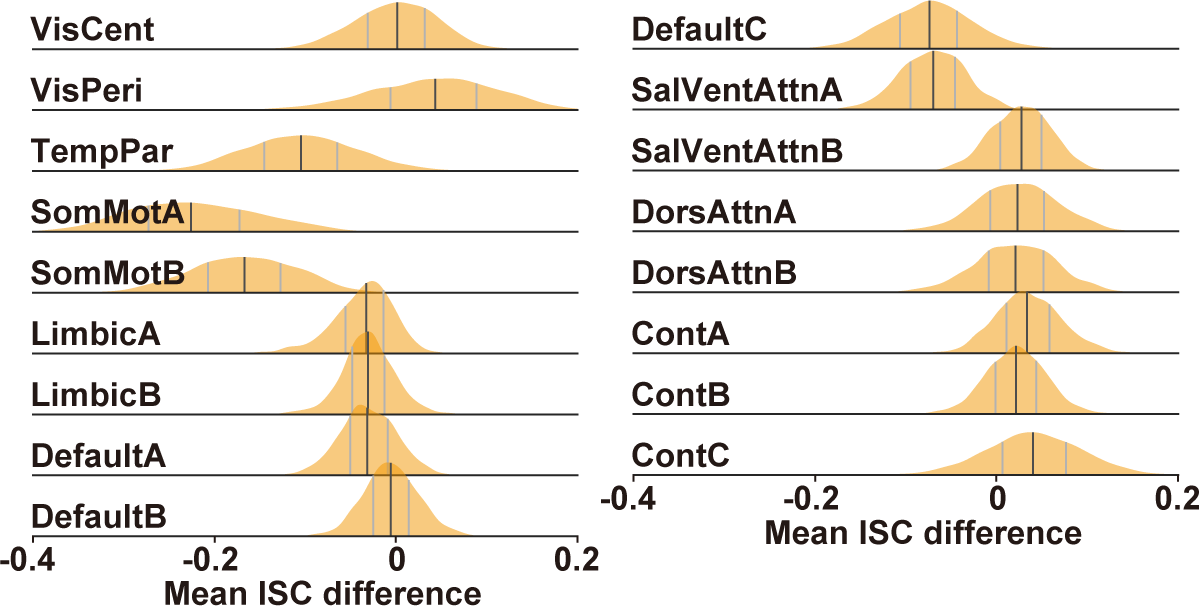
Distribution of ISC difference using the task-type model. Distributions of difference between the mean RSM- and Weight-ISC values (averaged across vertices) are averaged across vertices for the 17 network atlas (Schaefer et al. 2018), with the median (black line), 25 and 75 percentiles (gray lines).

**Figure S7.**
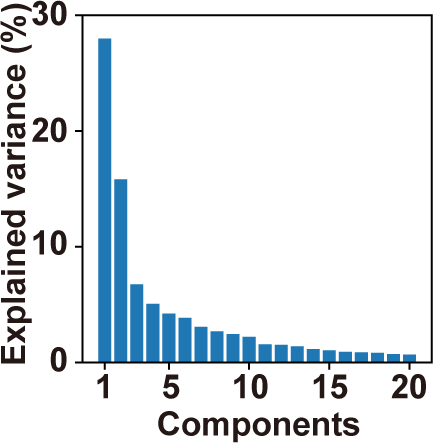
Explained variances of the top 20 principal components. The explained variances of the original weight matrix were obtained from principal component analysis (PCA) using half of the subjects (30 subjects; ID30-ID59), within a cortical mask of the RSM-vs. weight-ISC contrast with the other half of the subjects (29 subjects; ID01-ID29).

**Figure S8.**
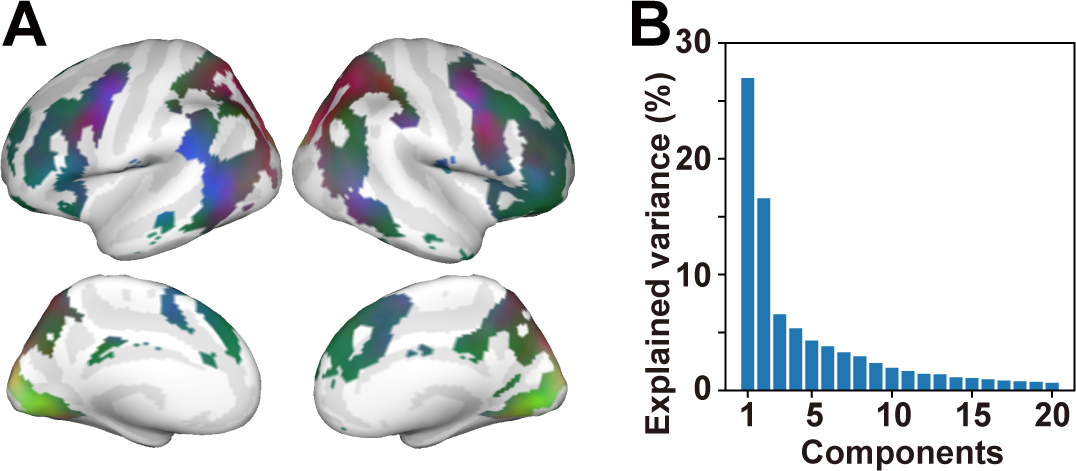
Principal component analysis (PCA) based on the other half of the subjects. **(A)** The PCA reveals a cortical map highlighting various RSM patterns. Vertices with a significant RSM-vs. weight-ISC value were masked using data from half of the subjects (subject ID30-ID59), and PCA was subsequently applied to the remaining subjects (ID01-ID29). Contributions of the top three PCs (PC1–PC3) are shown in red, green, and blue, respectively. **(B)** The explained variances of PCA are depicted for the top 20 components. The top three principal components account for 50.1% of the variances in the original weight matrices.

**Figure S9.**
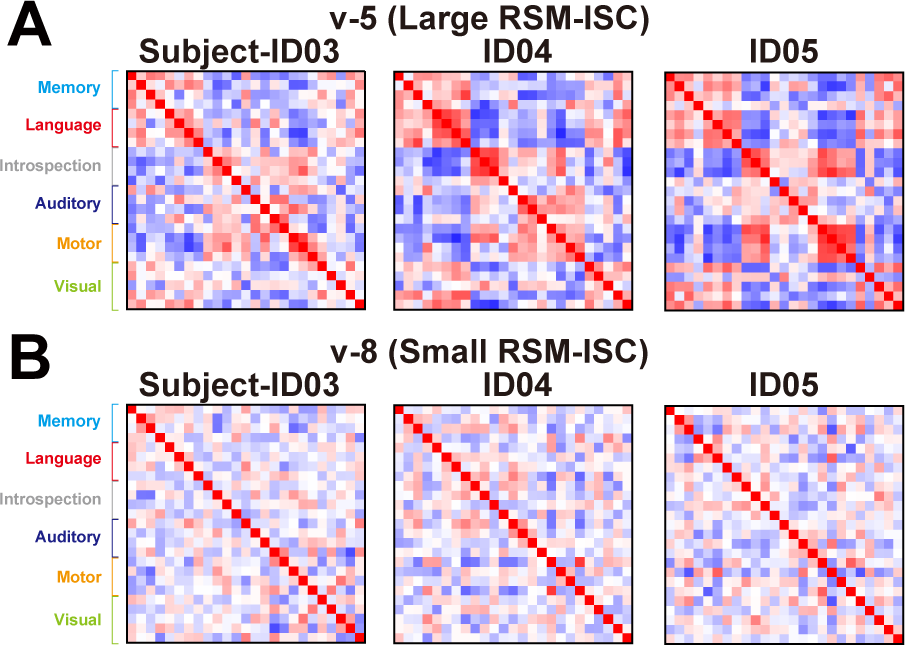
Individual variability in RSMs. **(A-B)** RSMs of three subjects (ID03, ID04, ID05) are displayed. These RSMs are derived from two example vertices (v-5, v-8) as referenced in Figure 4A. The RSMs are taken from vertices with **(A)** a large RSM-ISC and **(B)** a small RSM-ISC.

**Figure S10.**
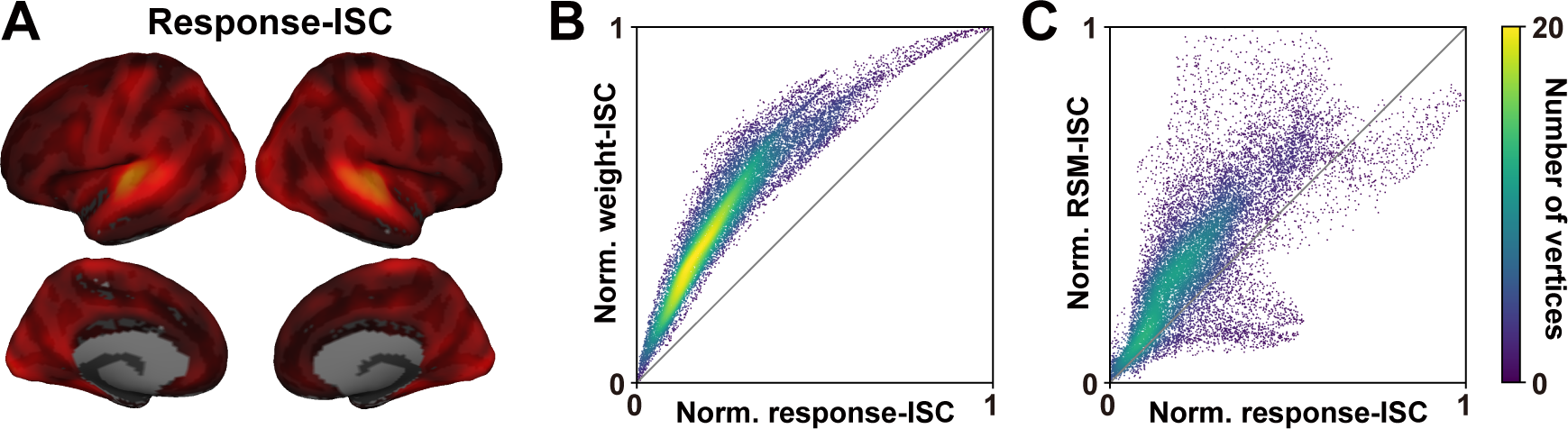
Comparison between model-free and model-based ISC. (A) The response-ISC is derived directly from brain responses, similar to the weight-ISC, and is visualized on the cortical surface. (B-C) Normalized scatter plots depict (B) the weight-ISC and (C) the RSM-ISC in relation to the response-ISC.

**Figure S11.**
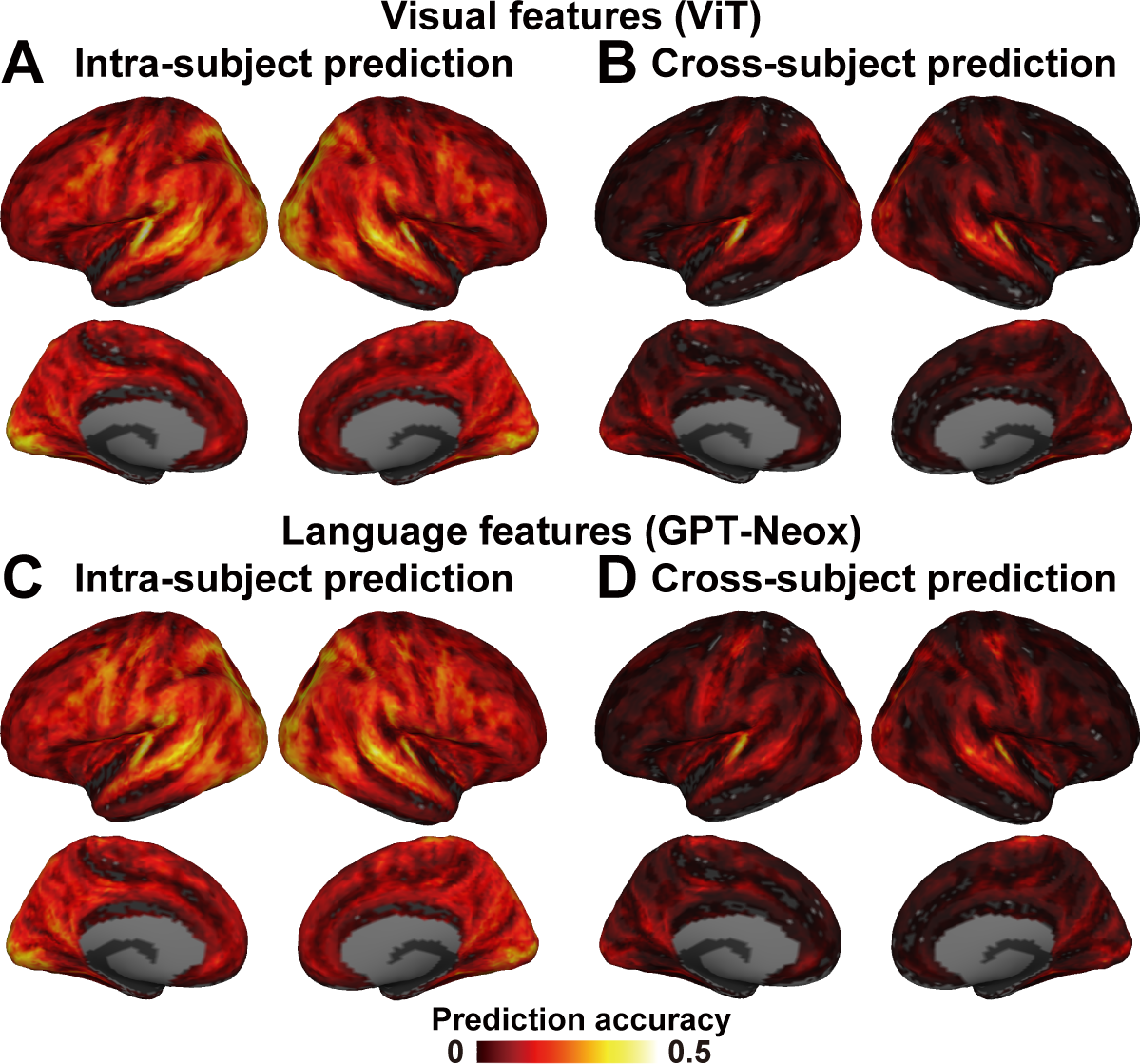
Prediction performance of latent features. (**A-D**) Prediction accuracies of encoding models utilizing latent visual features (ViT) and language features (GPT-Neox) are visualized on the standard cortical surface. The models are evaluated through **(A, C)** intra-subject prediction, using the same subject, and **(B, D)** cross-subject prediction, involving different subjects.

**Figure S12.**
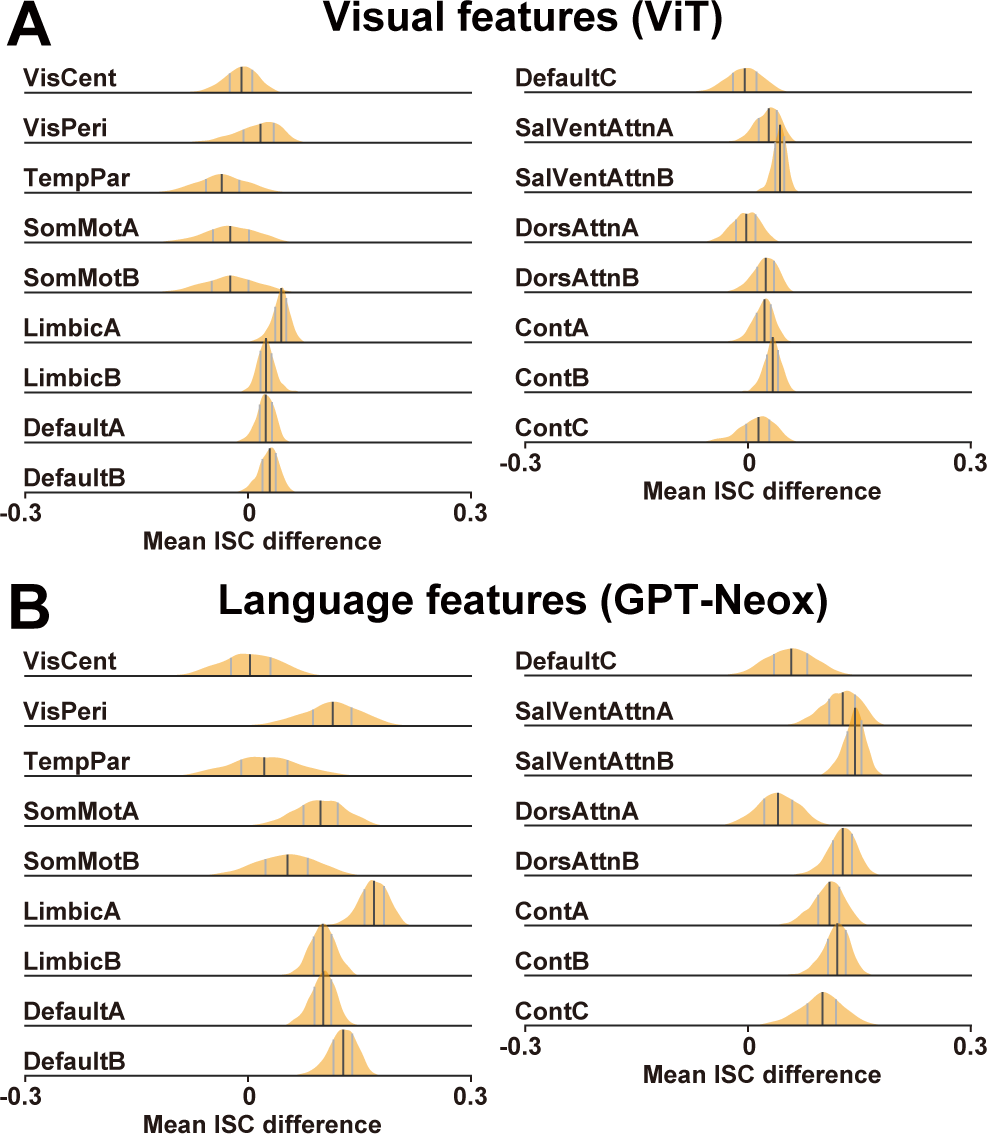
Distribution of the ISC difference using latent features. (**A-B**) Distributions of the difference between mean RSM- and weight-ISC values (averaged across vertices) are depicted for (**A**) visual and (**B**) language latent features across the 17 network atlases (Schaefer et al. 2018). The median is represented by the black line, and the 25 and 75 percentiles are indicated by gray lines.

## Description of each task

### Memory tasks

1. CalcHard

Subjects tackled a challenging arithmetic problem involving two-digit numbers. Duration: 8 s.

2. PropLogic

Subjects engaged with a syllogism based on prepositional logical relationships, determining the validity of the conclusion. Duration: 10 s.

3. MemoryLetter

Subjects memorized a sequence of letters. Duration: 6 s.

4. MatchLetter

Subjects assessed whether a presented series of letters matched the one shown previously (corresponding to the letters memorized in the MemoryLetter task). Duration: 6 s.

### Language tasks

5. Recipe

Subjects evaluated whether a given recipe matched the actual recipe of a specified dish. Duration: 6 s.

6. WordMeaning

Subjects determined whether the meaning of a presented word aligned with the sentence displayed above the word. Duration: 6 s.

7. Metaphor

Subjects read a metaphorical text and assessed whether the writer’s intention matched the meaning indicated above the text. Duration: 8 s.

8. RatePoem

Subjects read a poem and rated its quality. Duration: 10 s.

### Introspection tasks

9. RecallPast

Subjects recollected a past event. Duration: 8 s.

10. ImaginePlace

Subjects envisioned a specific place. Duration: 7 s.

11. ImagineMove

Subjects imagined their body in motion. Duration: 7 s.

12. CategoryFluency

Subjects recalled as many words as possible belonging to a given word category. Duration: 8 s.

### Auditory tasks

13. CountTone

Subjects counted the number of presented tones. Duration: 8 s.

14. MusicCategory

Subjects judged whether the genre of a piece of music matched the name displayed on the screen. Duration: 10 s.

15. DailySound

Subjects listened to the sound of a tool used daily and determined whether its name matched the one displayed above the photo. Duration: 6 s.

16. EmotionVoice

Subjects listened to a voice with a specific emotion and assessed whether it matched the emotion indicated above the photo. Duration: 6 s.

### Motor tasks

17. EyeBlink

Subjects blinked their eyes as many times as possible. Duration: 8 s.

18. PressLR

Subjects pressed buttons (with their right or left hand) as many times as possible. Duration: 8 s.

19. FootLR

Subjects moved their feet as much as possible without moving their head. Duration: 8 s.

20. Tongue

Subjects moved their tongue as much as possible. Duration: 6 s.

### Visual tasks

21. RateDeliciousPic

Subjects viewed a photo of food and rated its deliciousness. Duration: 6 s.

22. RateBeautyPic

Subjects observed a photo and rated its beauty. Duration: 6 s.

23. AnimalPhoto

Subjects viewed a photo of an animal and judged whether its name matched the one displayed above the photo. Duration: 6 s.

24. CountryMap

Subjects looked at a photo of a country map and determined whether its name (nation) matched the one displayed above the photo. Duration: 6 s.

25. ComparePeople

Subjects compared two photos of people and assessed whether they were the same person. Duration: 6 s.

## References

Abrams, Daniel A., Srikanth Ryali, Tianwen Chen, Parag Chordia, Amirah Khouzam, Daniel J. Levitin, and Vinod Menon. 2013. “Inter-Subject Synchronization of Brain Responses during Natural Music Listening.” The European Journal of Neuroscience 37 (9): 1458–69.

Avants, Brian B., Nicholas J. Tustison, Gang Song, Philip A. Cook, Arno Klein, and James C. Gee. 2011. “A Reproducible Evaluation of ANTs Similarity Metric Performance in Brain Image Registration.” NeuroImage 54 (3): 2033–44.

Benjamini, Yoav, and Yosef Hochberg. 1995. “Controlling the False Discovery Rate: A Practical and Powerful Approach to Multiple Testing.” Journal of the Royal Statistical Society 57 (1): 289–300.

Black, Sid, Stella Biderman, Eric Hallahan, Quentin Anthony, Leo Gao, Laurence Golding, Horace He, et al. 2022. “GPT-NeoX-20B: An Open-Source Autoregressive Language Model.” ArXiv [Cs.CL]. arXiv. http://arxiv.org/abs/2204.06745.

Brooks, Jeffrey A., Junichi Chikazoe, Norihiro Sadato, and Jonathan B. Freeman. 2019. “The Neural Representation of Facial-Emotion Categories Reflects Conceptual Structure.” Proceedings of the National Academy of Sciences of the United States of America 116 (32): 15861–70.

Carroll, J. Douglas, and Jih-Jie Chang. 1970. “Analysis of Individual Differences in Multidimensional Scaling via an N-Way Generalization of ‘Eckart-Young’ Decomposition.” Psychometrika 35 (3): 283–319.

Charest, Ian, Rogier A. Kievit, Taylor W. Schmitz, Diana Deca, and Nikolaus Kriegeskorte. 2014. “Unique Semantic Space in the Brain of Each Beholder Predicts Perceived Similarity.” Proceedings of the National Academy of Sciences of the United States of America 111 (40): 14565–70.

Chen, Janice, Yuan Chang Leong, Christopher J. Honey, Chung H. Yong, Kenneth A. Norman, and Uri Hasson. 2017. “Shared Memories Reveal Shared Structure in Neural Activity across Individuals.” Nature Neuroscience 20 (1): 115–25.

Cole, Michael W., Jeremy R. Reynolds, Jonathan D. Power, Grega Repovs, Alan Anticevic, and Todd S. Braver. 2013. “Multi-Task Connectivity Reveals Flexible Hubs for Adaptive Task Control.” Nature Neuroscience 16 (9): 1348–55.

Cui, Zaixu, Hongming Li, Cedric H. Xia, Bart Larsen, Azeez Adebimpe, Graham L. Baum, Matt Cieslak, et al. 2020. “Individual Variation in Functional Topography of Association Networks in Youth.” Neuron 106 (2): 340–353.e8.

Çukur, Tolga, Shinji Nishimoto, Alexander G. Huth, and Jack L. Gallant. 2013. “Attention during Natural Vision Warps Semantic Representation across the Human Brain.” Nature Neuroscience 16 (6): 763–70.

Cutts, Sarah A., Joshua Faskowitz, Richard F. Betzel, and Olaf Sporns. 2022. “Uncovering Individual Differences in Fine-Scale Dynamics of Functional Connectivity.” Cerebral Cortex, June. 10.1093/cercor/bhac214.

Dale, A. M., B. Fischl, and M. I. Sereno. 1999. “Cortical Surface-Based Analysis. I. Segmentation and Surface Reconstruction.” NeuroImage 9 (2): 179–94.

Devereux, Barry J., Alex Clarke, Andreas Marouchos, and Lorraine K. Tyler. 2013. “Representational Similarity Analysis Reveals Commonalities and Differences in the Semantic Processing of Words and Objects.” The Journal of Neuroscience: The Official Journal of the Society for Neuroscience 33 (48): 18906–16.

Dosovitskiy, A., L. Beyer, A. Kolesnikov, and D. Weissenborn. 2020. “Transformers for Image Recognition at Scale.” ArXiv Preprint ArXiv.

Dubois, Julien, and Ralph Adolphs. 2016. “Building a Science of Individual Differences from FMRI.” Trends in Cognitive Sciences 20 (6): 425–43.

Esteban, Oscar, Christopher J. Markiewicz, Ross W. Blair, Craig A. Moodie, A. Ilkay Isik, Asier Erramuzpe, James D. Kent, et al. 2019. “FMRIPrep: A Robust Preprocessing Pipeline for Functional MRI.” Nature Methods 16 (1): 111–16.

Feilong, Ma, Samuel A. Nastase, J. Swaroop Guntupalli, and James V. Haxby. 2018. “Reliable Individual Differences in Fine-Grained Cortical Functional Architecture.” NeuroImage 183 (December): 375–86.

Finn, Emily S., Enrico Glerean, Arman Y. Khojandi, Dylan Nielson, Peter J. Molfese, Daniel A. Handwerker, and Peter A. Bandettini. 2020. “Idiosynchrony: From Shared Responses to Individual Differences during Naturalistic Neuroimaging.” NeuroImage 215 (July): 116828.

Friedman, Naomi P., and Trevor W. Robbins. 2022. “The Role of Prefrontal Cortex in Cognitive Control and Executive Function.” Neuropsychopharmacology: Official Publication of the American College of Neuropsychopharmacology 47 (1): 72–89.

Gao, James S., Alexander G. Huth, Mark D. Lescroart, and Jack L. Gallant. 2015. “Pycortex: An Interactive Surface Visualizer for FMRI.” Frontiers in Neuroinformatics 9 (September): 23.

Gordon, Evan M., Timothy O. Laumann, Adrian W. Gilmore, Dillan J. Newbold, Deanna J. Greene, Jeffrey J. Berg, Mario Ortega, et al. 2017. “Precision Functional Mapping of Individual Human Brains.” Neuron 95 (4): 791–807.e7.

Gratton, Caterina, Timothy O. Laumann, Ashley N. Nielsen, Deanna J. Greene, Evan M. Gordon, Adrian W. Gilmore, Steven M. Nelson, et al. 2018. “Functional Brain Networks Are Dominated by Stable Group and Individual Factors, Not Cognitive or Daily Variation.” Neuron 98 (2): 439–452.e5.

Greve, Douglas N., and Bruce Fischl. 2009. “Accurate and Robust Brain Image Alignment Using Boundary-Based Registration.” NeuroImage 48 (1): 63–72.

Guntupalli, J. Swaroop, Ma Feilong, and James V. Haxby. 2018. “A Computational Model of Shared Fine-Scale Structure in the Human Connectome.” PLoS Computational Biology 14 (4): e1006120.

Guntupalli, J. Swaroop, Michael Hanke, Yaroslav O. Halchenko, Andrew C. Connolly, Peter J. Ramadge, and James V. Haxby. 2016. “A Model of Representational Spaces in Human Cortex.” Cerebral Cortex 26 (6): 2919–34.

Hasson, Uri, Orit Furman, Dav Clark, Yadin Dudai, and Lila Davachi. 2008. “Enhanced Intersubject Correlations during Movie Viewing Correlate with Successful Episodic Encoding.” Neuron 57 (3): 452–62.

Hasson, Uri, Yuval Nir, Ifat Levy, Galit Fuhrmann, and Rafael Malach. 2004. “Intersubject Synchronization of Cortical Activity during Natural Vision.” Science 303 (5664): 1634–40.

Horikawa, Tomoyasu, Alan S. Cowen, Dacher Keltner, and Yukiyasu Kamitani. 2020. “The Neural Representation of Visually Evoked Emotion Is High-Dimensional, Categorical, and Distributed across Transmodal Brain Regions.” IScience 23 (5): 101060.

Horikawa, Tomoyasu, and Yukiyasu Kamitani. 2017. “Generic Decoding of Seen and Imagined Objects Using Hierarchical Visual Features.” Nature Communications 8 (May): 15037.

Huth, Alexander G., Wendy A. de Heer, Thomas L. Griffiths, Frédéric E. Theunissen, and Jack L. Gallant. 2016. “Natural Speech Reveals the Semantic Maps That Tile Human Cerebral Cortex.” Nature 532 (7600): 453–58.

Ito, Takuya, and John D. Murray. 2022. “Multitask Representations in the Human Cortex Transform along a Sensory-to-Motor Hierarchy.” Nature Neuroscience, December. 10.1038/s41593-022-01224-0.

Jenkinson, Mark, Peter Bannister, Michael Brady, and Stephen Smith. 2002. “Improved Optimization for the Robust and Accurate Linear Registration and Motion Correction of Brain Images.” NeuroImage 17 (2): 825–41.

Kauppi, Jukka-Pekka, Juha Pajula, Jari Niemi, Riitta Hari, and Jussi Tohka. 2017. “Functional Brain Segmentation Using Inter-Subject Correlation in FMRI.” Human Brain Mapping 38 (5): 2643–65.

Kay, Kendrick N., Stephen V. David, Ryan J. Prenger, Kathleen A. Hansen, and Jack L. Gallant. 2008. “Modeling Low-Frequency Fluctuation and Hemodynamic Response Timecourse in Event-Related FMRI.” Human Brain Mapping 29 (2): 142–56.

Kay, Kendrick N., Thomas Naselaris, Ryan J. Prenger, and Jack L. Gallant. 2008. “Identifying Natural Images from Human Brain Activity.” Nature 452 (7185): 352–55.

Keller, Arielle S., Valerie J. Sydnor, Adam Pines, Damien A. Fair, Dani S. Bassett, and Theodore D. Satterthwaite. 2023. “Hierarchical Functional System Development Supports Executive Function.” Trends in Cognitive Sciences 27 (2): 160–74.

Koide-Majima, Naoko, Tomoya Nakai, and Shinji Nishimoto. 2020. “Distinct Dimensions of Emotion in the Human Brain and Their Representation on the Cortical Surface.” NeuroImage 222 (November): 117258.

Kriegeskorte, Nikolaus, Rainer Goebel, and Peter Bandettini. 2006. “Information-Based Functional Brain Mapping.” Proceedings of the National Academy of Sciences of the United States of America 103 (10): 3863–68.

Kriegeskorte, Nikolaus, and Rogier A. Kievit. 2013. “Representational Geometry: Integrating Cognition, Computation, and the Brain.” Trends in Cognitive Sciences 17 (8): 401–12.

Leeuw, Jan de, and Patrick Mair. 2009. “Multidimensional Scaling Using Majorization: SMACOF in R.” Journal of Statistical Software 31 (August): 1–30.

Margulies, Daniel S., Satrajit S. Ghosh, Alexandros Goulas, Marcel Falkiewicz, Julia M. Huntenburg, Georg Langs, Gleb Bezgin, et al. 2016. “Situating the Default-Mode Network along a Principal Gradient of Macroscale Cortical Organization.” Proceedings of the National Academy of Sciences of the United States of America 113 (44): 12574–79.

Michon, Katherine J., Dalia Khammash, Molly Simmonite, Abbey M. Hamlin, and Thad A. Polk. 2022. “Person-Specific and Precision Neuroimaging: Current Methods and Future Directions.” NeuroImage, August, 119589.

Mueller, Sophia, Danhong Wang, Michael D. Fox, B. T. Thomas Yeo, Jorge Sepulcre, Mert R. Sabuncu, Rebecca Shafee, Jie Lu, and Hesheng Liu. 2013. “Individual Variability in Functional Connectivity Architecture of the Human Brain.” Neuron 77 (3): 586–95.

Nakai, Tomoya, Naoko Koide-Majima, and Shinji Nishimoto. 2021. “Correspondence of Categorical and Feature-Based Representations of Music in the Human Brain.” Brain and Behavior 11 (1): e01936.

Nakai, Tomoya, and Shinji Nishimoto. 2020. “Quantitative Models Reveal the Organization of Diverse Cognitive Functions in the Brain.” Nature Communications 11 (1): 1142.

Nakai, Tomoya, and Shinji Nishimoto. 2022. “Representations and Decodability of Diverse Cognitive Functions Are Preserved across the Human Cortex, Cerebellum, and Subcortex.” Communications Biology 5 (1): 1245.

Nakai, Tomoya, and Shinji Nishimoto. 2023a. “Quantitative Modelling Demonstrates Format-Invariant Representations of Mathematical Problems in the Brain.” The European Journal of Neuroscience, January. 10.1111/ejn.15925.

Nakai, Tomoya, and Shinji Nishimoto. 2023b. “Artificial Neural Network Modelling of the Neural Population Code Underlying Mathematical Operations.” NeuroImage, February, 119980.

Nakai, Tomoya, Hiroto Q. Yamaguchi, and Shinji Nishimoto. 2021. “Convergence of Modality Invariance and Attention Selectivity in the Cortical Semantic Circuit.” Cerebral Cortex 31 (10): 4825–39.

Naselaris, Thomas, Kendrick N. Kay, Shinji Nishimoto, and Jack L. Gallant. 2011. “Encoding and Decoding in FMRI.” NeuroImage 56 (2): 400–410.

Nastase, Samuel A., Valeria Gazzola, Uri Hasson, and Christian Keysers. 2019. “Measuring Shared Responses across Subjects Using Intersubject Correlation.” Social Cognitive and Affective Neuroscience 14 (6): 667–85.

Niendam, Tara A., Angela R. Laird, Kimberly L. Ray, Y. Monica Dean, David C. Glahn, and Cameron S. Carter. 2012. “Meta-Analytic Evidence for a Superordinate Cognitive Control Network Subserving Diverse Executive Functions.” Cognitive, Affective & Behavioral Neuroscience 12 (2): 241–68.

Nishida, Satoshi, and Shinji Nishimoto. 2018. “Decoding Naturalistic Experiences from Human Brain Activity via Distributed Representations of Words.” NeuroImage 180 (Pt A): 232–42.

Nishimoto, Shinji, An T. Vu, Thomas Naselaris, Yuval Benjamini, Bin Yu, and Jack L. Gallant. 2011. “Reconstructing Visual Experiences from Brain Activity Evoked by Natural Movies.” Current Biology: CB 21 (19): 1641–46.

Norman-Haignere, Sam, Nancy G. Kanwisher, and Josh H. McDermott. 2015. “Distinct Cortical Pathways for Music and Speech Revealed by Hypothesis-Free Voxel Decomposition.” Neuron 88 (6): 1281–96.

Peelen, Marius V., and Paul E. Downing. 2023. “Testing Cognitive Theories with Multivariate Pattern Analysis of Neuroimaging Data.” Nature Human Behaviour, August. 10.1038/s41562-023-01680-z.

Popham, Sara F., Alexander G. Huth, Natalia Y. Bilenko, Fatma Deniz, James S. Gao, Anwar O. Nunez-Elizalde, and Jack L. Gallant. 2021. “Visual and Linguistic Semantic Representations Are Aligned at the Border of Human Visual Cortex.” Nature Neuroscience 24 (11): 1628–36.

Power, Jonathan D., Bradley L. Schlaggar, Christina N. Lessov-Schlaggar, and Steven E. Petersen. 2013. “Evidence for Hubs in Human Functional Brain Networks.” Neuron 79 (4): 798–813.

Ren, Yudan, Vinh Thai Nguyen, Lei Guo, and Christine Cong Guo. 2017. “Inter-Subject Functional Correlation Reveal a Hierarchical Organization of Extrinsic and Intrinsic Systems in the Brain.” Scientific Reports 7 (1): 10876.

Schaefer, Alexander, Ru Kong, Evan M. Gordon, Timothy O. Laumann, Xi-Nian Zuo, Avram J. Holmes, Simon B. Eickhoff, and B. T. Thomas Yeo. 2018. “Local-Global Parcellation of the Human Cerebral Cortex from Intrinsic Functional Connectivity MRI.” Cerebral Cortex 28 (9): 3095–3114.

Schrimpf, Martin, Idan Asher Blank, Greta Tuckute, Carina Kauf, Eghbal A. Hosseini, Nancy Kanwisher, Joshua B. Tenenbaum, and Evelina Fedorenko. 2021. “The Neural Architecture of Language: Integrative Modeling Converges on Predictive Processing.” Proceedings of the National Academy of Sciences of the United States of America 118 (45). 10.1073/pnas.2105646118.

Seghier, Mohamed L., and Cathy J. Price. 2018. “Interpreting and Utilising Intersubject Variability in Brain Function.” Trends in Cognitive Sciences 22 (6): 517–30.

Seitzman, Benjamin A., Caterina Gratton, Timothy O. Laumann, Evan M. Gordon, Babatunde Adeyemo, Ally Dworetsky, Brian T. Kraus, et al. 2019. “Trait-like Variants in Human Functional Brain Networks.” Proceedings of the National Academy of Sciences of the United States of America 116 (45): 22851–61.

Setti, Francesca, Giacomo Handjaras, Davide Bottari, Andrea Leo, Matteo Diano, Valentina Bruno, Carla Tinti, et al. 2023. “A Modality-Independent Proto-Organization of Human Multisensory Areas.” Nature Human Behaviour, January. 10.1038/s41562-022-01507-3.

Sheng, Jintao, Sisi Wang, Liang Zhang, Chuqi Liu, Liang Shi, Yu Zhou, Huinan Hu, Chuansheng Chen, and Gui Xue. 2023. “Intersubject Similarity in Neural Representations Underlies Shared Episodic Memory Content.” Proceedings of the National Academy of Sciences of the United States of America 120 (35): e2308951120.

Silbert, Lauren J., Christopher J. Honey, Erez Simony, David Poeppel, and Uri Hasson. 2014. “Coupled Neural Systems Underlie the Production and Comprehension of Naturalistic Narrative Speech.” Proceedings of the National Academy of Sciences of the United States of America 111 (43): E4687–96.

Simony, Erez, Christopher J. Honey, Janice Chen, Olga Lositsky, Yaara Yeshurun, Ami Wiesel, and Uri Hasson. 2016. “Dynamic Reconfiguration of the Default Mode Network during Narrative Comprehension.” Nature Communications 7 (July): 12141.

Stoecklein, Sophia, Anne Hilgendorff, Meiling Li, Kai Förster, Andreas W. Flemmer, Franziska Galiè, Stephan Wunderlich, et al. 2020. “Variable Functional Connectivity Architecture of the Preterm Human Brain: Impact of Developmental Cortical Expansion and Maturation.” Proceedings of the National Academy of Sciences of the United States of America 117 (2): 1201–6.

Vanderwal, Tamara, Jeffrey Eilbott, Emily S. Finn, R. Cameron Craddock, Adam Turnbull, and F. Xavier Castellanos. 2017. “Individual Differences in Functional Connectivity during Naturalistic Viewing Conditions.” NeuroImage 157 (August): 521–30.

Wang, Danhong, Randy L. Buckner, Michael D. Fox, Daphne J. Holt, Avram J. Holmes, Sophia Stoecklein, Georg Langs, et al. 2015. “Parcellating Cortical Functional Networks in Individuals.” Nature Neuroscience 18 (12): 1853–60.

Wilson, Stephen M., Istvan Molnar-Szakacs, and Marco Iacoboni. 2008. “Beyond Superior Temporal Cortex: Intersubject Correlations in Narrative Speech Comprehension.” Cerebral Cortex 18 (1): 230–42.

Zhang, Y., M. Brady, and S. Smith. 2001. “Segmentation of Brain MR Images through a Hidden Markov Random Field Model and the Expectation-Maximization Algorithm.” IEEE Transactions on Medical Imaging 20 (1): 45–57.

